# A Solanoeclepin A precursor functions as a new rhizosphere signaling molecule recruiting growth-promoting microbes under nitrogen deficiency

**DOI:** 10.64898/2025.12.29.696744

**Authors:** Davar Abedini, Alessandra Guerrieri, Rahul Jain, Fred White, Joep Koomen, Yuting Yang, Kangning Wang, Gertjan Kramer, Harro Bouwmeester, Lemeng Dong

**Affiliations:** Plant Hormone Biology Group, Swammerdam Institute for Life Sciences (SILS), University of Amsterdam, Amsterdam, The Netherlands; Biosystems Data Analysis Group, Swammerdam Institute for Life Sciences, University of Amsterdam, The Netherlands; Laboratory for Mass Spectrometry of Biomolecules, Swammerdam Institute for Life Sciences (SILS), University of Amsterdam, Amsterdam, The Netherlands

**Keywords:** Solanoeclepin A, Cyst Nematodes, Nitrogen Deficiency, Triterpenes, Rhizosphere Beneficial Microbes, Signaling Molecule

## Abstract

Solanoeclepin A (SolA) is a triterpenoid exuded from the roots of Solanaceae plants, originally identified as hatching stimulant for plant parasitic cyst nematodes. Assuming that evolution would have selected against the production of such a fitness lowering molecule, we postulated that SolA must serve another, beneficial, role for the plant. In this study, we demonstrate that nitrogen (N) deficiency strongly increases the SolA concentration in tomato root exudate. Moreover, SolA is produced only under non-sterile conditions, indicating that soil microbiota are involved in its production. Time-resolved RNAseq analysis revealed several candidate genes for SolA biosynthesis, which were all upregulated under N deficiency. Transient silencing of two SolA biosynthetic genes (e.g., *CYP749A19* and *CYP749A20*) significantly reduced SolA production. Microbiome analysis on the rhizosphere of these plants demonstrated that the recruitment of beneficial *Massilia* spp. was inhibited in the transiently silenced plants. Isolation of a *Massilia* strain, identified as *Massilia cellulosiltytica*, allowed us to show that it has growth-promoting activity under N deficiency, likely via indole-3-acetic acid production and enhanced N acquisition. Root exudate of N-starved tomato displayed strong chemotactic activity towards this strain. Together, these findings demonstrate that SolA production under N deficiency relies on the interaction between tomato and soil microbiota that convert a plant-produced precursor that is a recruitment signal for beneficial, growth-promoting microbes to SolA, which was hijacked by cyst nematodes as a reliable host presence cue. This dual functionality resembles the microbial transformation of primary into secondary bile acids, important signalling molecules, in the gut of animals, and suggests convergent evolution in host–microbe co-metabolism. Overall, our study positions SolA as a multifunctional signaling molecule in rhizosphere interactions, with potential application for enhancing crop resilience and sustainable pest management.

## Introduction

Agriculture consumes large amounts of non-renewable resources and is one of the major contributors to environmental pollution and climate change (Filonchyk et al., 2024). Chemical pesticides accumulate in soils and waterways, while fertiliser runoff causes eutrophication (Akinnawo, 2023). Fertiliser use also generates CO_2_ and N_2_O emissions, with nitrogen fertilizer production relying on the energy-intensive Haber-Bosch process (Jacoby et al., 2017), and microbial activity releasing greenhouse gasses, NH_3_ and N_2_O (Munch and Velthof, 2007). Phosphorus fertiliser, derived from finite rock phosphate, may be depleted in 100 years (Fayiga and Nwoke, 2016). Pesticide use has doubled since 1990, reaching 3.7 million tonnes in 2022 (FAO, 2024).

Solutions for these enormous challenges may be found in the rhizosphere where there is growing evidence that microbiota can potentially alleviate the need for pesticide and fertiliser use in agriculture. The rhizosphere is heavily influenced by plants through their root exudate, which plays a role in recruiting beneficial microbes and deterring pathogens. Through their root exudate, plants exude 5-30% of photosynthetically fixed carbon into the rhizosphere (Sasse et al., 2018) to foster interactions with root microbiota. One of the classes of metabolites that are present in plant root exudate are the triterpenes. They originate from the 30-carbon precursor squalene and represent a vast and varied class of natural products with over 20,000 identified compounds, primarily in the plant kingdom (Thimmappa et al., 2014; Jia Liu et al., 2024). Encompassing various subclasses such as sterols and saponins, these specialized metabolites have gained significant attention due to their multifaceted roles in plant physiology and ecology (Holopainen et al., 2013; Cárdenas et al., 2019). For instance, β-amyrin is involved in root development in oat (Kemen et al., 2014) and *Lotus japonicus* (Krokida et al., 2013), but also is a precursor for complex triterpene glycosides associated with plant defense (Lee et al., 2001; Kemen et al., 2014). Triterpenes have also been shown to play a role in shaping the rhizosphere microbiome, influencing the composition and diversity of root-associated microorganisms. Cucurbitacins, for example, originally identified as plant defense compounds against pests and pathogens (Dinan, 1997), affect the root microbiome in cucumber (Zhung et al., 2022). Also in *Arabidopsis thaliana,* two tricyclic triterpene esters, arabidin and thalian, which are exuded into the rhizosphere, were demonstrated to affect the root microbiome (Huang et al., 2019).

The biosynthesis of triterpenoids and its regulation are intricate processes fine-tuned by both internal signals and environmental factors including biotic and abiotic stresses. Typically, abscisic acid (ABA) regulates the response against abiotic stresses such as salinity, drought, cold and heat stress (Verma et al., 2016) while salicylic acid (SA) and jasmonate (JA) play major roles in the response to biotic stress conditions such as pathogen infection and insect attack. Most triterpenoids were shown to be induced by jasmonate, for example, in *Medicago truncatula* (Mertens et al., 2016), *Centella asiatica* (Mangas et al., 2006), and sweet basil (Misra et al., 2014). However, drought stress and ABA induce the production of the triterpenoid cucurbitacin C in cucumber (Shang et al., 2014), while nitrogen starvation negatively affects triterpenoid withanolide biosynthesis in *Withania somnifera* (Devkar et al., 2015). The changes in triterpenoid production mentioned here often strongly correlate with the expression of the corresponding biosynthetic pathway genes (Kutschera 2012; Merten 2016; Misra 2014; Shang 2014).

Solanoeclepin A (SolA) and glycinoeclepins form a sub-class of highly bioactive and oxygenated nortriterpenoids that were discovered decades ago for their hatching stimulant activity in plant-parasitic cyst nematodes (CNs) that affect crops such as potato and kidney bean, respectively (Masamune et al., 1982; Fukuzawa et al., 1985; Mulder et al., 1996). SolA has been reported to be produced in extremely low concentrations in the root exudates of potato and tomato (Guerrieri et al., 2021; Vlaar, et al., 2022).

The production of these hatching stimulants by plants forms a conundrum: considering that they negatively impact plant fitness by inducing hatching in pathogenic cyst nematodes, evolution should have selected against the production of these molecules. Unless we are overlooking a positive role for these compounds with a fitness benefit that compensates the negative impact of hatching. This is reminiscent of the strigolactones, which were originally discovered as germination stimulants of root parasitic plants and later demonstrated to be a recruitment signal for arbuscular mycorrhizal fungi and other potentially beneficial micro-organisms (Akiyama and Hayashi, 2006; Kim et al., 2022). Here we postulate that this must indeed be the case: the eclepins have a beneficial role, which we have overlooked so far. In the present study we set out to discover that role.

A beneficial role for the eclepins, alongside their negative activity as hatching stimulants, could impose both positive and negative selection pressures. This combination may drive structural diversification, resulting in an evolutionary arms race, as has been extensively postulated for the strigolactones. (Wang and Bouwmeester, 2018; Bouwmeester et al., 2021). Indeed, in kidney bean, three different eclepins were identified, glycinoeclepin A, B and C (Masamune et al., 1982; Fukuzawa et al., 1985), and recently also in tomato a second and third solanoeclepin - both unfortunately called solanoeclepin B although they seem to have a different structure - were reported (Vlaar et al., 2022a; Shimizu et al., 2023). Shimizu *et al*. (2023) proposed that Solanoeclepin B (SolB) is the plant-exuded precursor of SolA. However, Akiyama *et al*., (2025) later identified Solanoeclepin C (SolC), an acetylated derivative of SolB, and showed it is secreted by tomato roots and microbially converted to SolB, which is subsequently converted to SolA.

Recently, Shimizu and colleagues (2023) reported several biosynthetic genes, including cytochrome P450 monooxygenases (CYPs) and 2-oxoglutarate-dependent dioxygenases (DOXs), as being involved in SolA production, or rather in the production of SolC, given that SolC is the eclepin exuded by roots.

To get more insight into the biological significance of the eclepins, in the present study we investigated the effect of environmental factors on SolA biosynthesis. This demonstrated that it is strongly upregulated under N deficiency. This allowed us to use a gene cloning strategy in which we searched for genes whose expression is also upregulated by N deficiency, in a pattern similar to the upregulation of SolA biosynthesis. This experiment also showed that the presence of microbiota is required for the production of SolA, from a precursor that is exuded by tomato roots, under N deficiency. Through transient silencing of two SolA biosynthetic genes, we demonstrated that they are indeed involved in the biosynthesis of SolA or rather its precursor. These transiently silenced plants allowed us to demonstrate that the biosynthesis of the SolA precursor results in the recruitment of various bacterial taxa, specifically beneficial *Massilia spp.*, which mitigate the effects of N deficiency and promote growth of tomato under N-deficient conditions.

## Results

### Nitrogen deficiency boosts SolA concentration in tomato root exudate

To identify conditions affecting SolA production, potato and tomato plants were subjected to a range of environmental factors, including variations in temperature, hormone treatments (including salicylic acid, and jasmonic acid), and nutrient availability including nitrogen (N) and phosphorous deficiencies. From all these treatments, N deficiency emerged as a critical factor boosting SolA production (Figure S1). To further explore the impact of N on SolA production, tomato plants were cultivated in an aeroponics system, allowing precise control over the nutrient concentration in the nutrient solution, as well as easy collection of root exudates (Figure 1A). Two weeks of N deficiency resulted in a significantly higher SolA accumulation in root exudate of tomato plants as compared to N-sufficient or increased N conditions (Figure S2). Interestingly, when N was reintroduced to N-starved plants, SolA levels significantly decreased (Figure 1B). Moreover, a PCN hatching assay using root exudates from N-starved and control (N-sufficient) tomato plants showed that exudate from N-starved plants induced significantly higher *Globodera pallida* hatching than the exudate of control plants grown under normal N (Figure 1C). Additionally, different N sources were tested, revealing that plants grown under nitrate-free conditions produced more SolA than those deprived of ammonia. However, complete N starvation had the strongest effect (Figure S3). These results suggest that N plays a crucial role in modulating SolA production.

**Figure 1.**
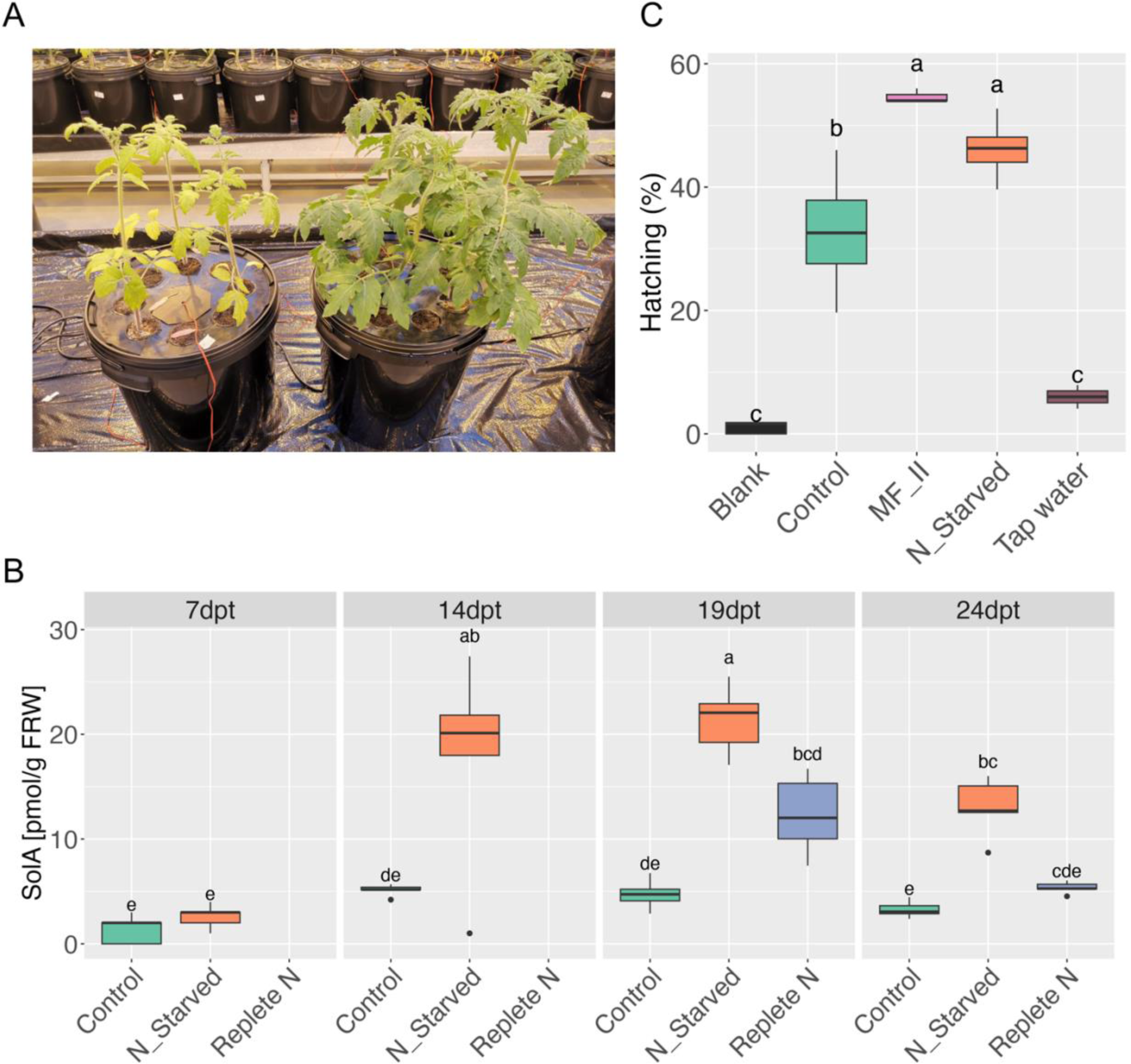
Nitrogen deficiency increases the SolA concentration in tomato root exudate. A) tomato plants growing on an aeroponics system with (Control) and without (N_Starved) nitrogen for 12 days after treatment. B) SolA concentration in the root exudate at 7-, 14-, 19-, and 24-days post-treatment (dpt) in nitrogen sufficient (Control), nitrogen starved (N_Starved), and recovered plants (Replete N) (N-starved for two weeks before nitrogen was repleted). C) PCN hatching assay with root exudate of plants grown without nitrogen (N_Starved) or with normal nitrogen (Control). As positive control the root exudate of potato variety MFII was used, which induces high PCN hatching. Hoagland solution (Blank) and tap water were used as negative controls.

### The presence of soil microbiota is essential for SolA formation

Tomato plants in the aeroponics system were grown either with a small amount of potting compost or in a rockwool cube. Interestingly, when grown on rockwool, N-deficient tomato plants did not produce SolA, whereas those grown in soil produced high levels of SolA (Figure 2A, Figure S4A). To find out whether this was due to the presence of microbiota in the potting compost we used sterilization of the soil through gamma-irradiation and autoclaving. Indeed, both treatments also prevented SolA production, just as the use of rockwool (Figure 2A). All this suggests that soil microbiota play a crucial, so far overlooked, role in SolA production.

**Figure 2.**
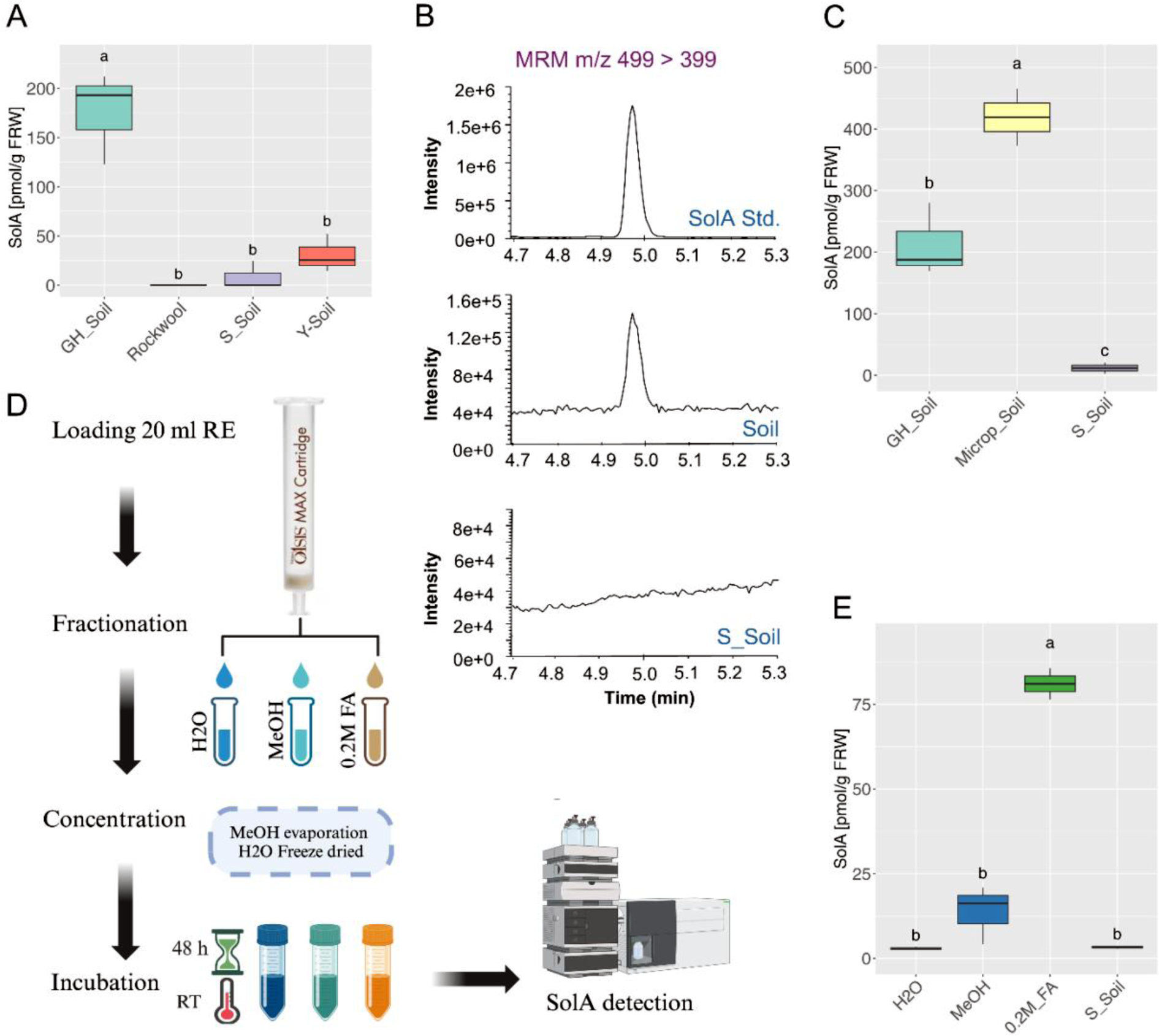
Microbial involvement in converting a SolA precursor to SolA. A) SolA concentration in root exudate of tomato seedlings grown in the aeroponics system on non-sterilized greenhouse soil (GH_Soil), autoclaved soil (S_Soil), gamma-irradiated soil (Y-Soil) and rockwool. B) Chromatograms demonstrating the SolA peak at *m/z* 499 > 399 in the standard (SoA Std.), non-sterilized soil (Soil), and autoclaved soil (S_Soil). C) Box plot showing the impact of soil type (GH_Soil, Greenhouse soil; MiCrop_Soil collected from Dutch fields; and S_Soil, autoclaved MiCrop soil) on the conversion of the SolA precursor to SolA. D) Ion exchange chromatography workflow for the fractionation and identification of SolA precursor(s) from tomato root exudate. 20 mL root exudate of N starved tomato plants growing on rockwool was fractionated using an Oasis MAX cartridge through sequential elution with water (H_2_O), methanol (MeOH), and 0.2 M formic acid in methanol (0.2M FA). After methanol evaporation and water removal through freeze-drying, the collected fractions were incubated with soil for 48 hours at room temperature. E) SolA was then extracted and quantified via UPLC MS/MS from these fractions.

These results led us to hypothesize that SolA is a molecule jointly synthesized by the plant and one or more soil microorganisms. In other words, we postulated that the plant produces a precursor that is released into the rhizosphere, where microbial activity converts it into SolA. To test this hypothesis, we developed a precursor bioconversion assay. In this assay we used root exudate of tomato plants grown under N deficiency on rockwool and incubated that with autoclaved (sterilized) or unsterilized soil (Figure S4B). When this root exudate, likely containing the SolA precursor, was incubated with unsterilized soil for 2 days, SolA formation occurred, while no such conversion was observed in sterilized soil (Figure 2B).

Incubation of root exudate with different soil types including potting soil and a field soil revealed substantial variation in the efficiency of this conversion process (Figure 2C), suggesting that the specific microbial community that is present in a certain soil contributes to SolA formation. These findings suggest that the presence of (certain) microbiota in the soil is essential for the conversion of a SolA precursor, present in the root exudate of tomato, into SolA.

To investigate the presence and characteristics of the SolA precursor in tomato root exudate, root exudate was fractionated using ion exchange chromatography on a MAX cartridge (Figure 2D). Elution was performed in three steps: first with water, then with methanol, and finally with 0.2 M formic acid in methanol. Each fraction was incubated with soil for 48 hours, after which SolA was extracted using solid-phase extraction (SPE). The results revealed that the fraction retained in the MAX column and eluted with 0.2 M formic acid in methanol contained the SolA precursor, as confirmed by the detection of SolA after incubation (Figure 2E). This suggests that, like SolA, its precursor likely already contains an acidic functional group, as it was retained in the ion exchange columns and only eluted with 0.2 M formic acid in methanol.

To investigate the presence of other SolA precursors, samples from the conversion assay were analyzed using quadrupole time-of-flight mass spectrometry (LC-qTOF-MS). Metabolic features were selected based on their *m/z* value, putative chemical formula, and relationship to the SolA biosynthesis pathway (details in Materials and Methods) (Table S1). C_26_H_28_O_9_, C_27_H_30_O_9_ (SolA), and C_27_H_32_O_9_ (Solanoeclepin B, SolB) (Shimizu et al., 2023) are virtually absent in a sterile soil conversion assay but significantly enriched in a conversion assay with non-sterile soil (19-fold, P

= 0.0015; 4-fold, P = 0.0039; 5.27-fold, P = 0.0487, respectively), supporting the assumption that these metabolites are products of microbial conversion. In contrast, C_30_H_46_O_6_ is highly abundant in the sterile soil, but significantly reduced in the non-sterile soil conversion assay (ratio = 0.13, P = 0.0117), suggesting microbial degradation or further transformation. Additionally, C_26_H_30_O_9_, C_26_H_32_O_9_, C_26_H_34_O_7_, C_27_H_36_O_7_, C_26_H_32_O_7_, C_26_H_32_O_8_, C_28_H_36_O_8_, and C_27_H_34_O_7_ were detected in small amounts in the sterile soil while they slightly, but non-significantly, increased in the non-sterile soil conversion assay. This suggests that these compounds might serve as intermediate metabolites in microbial conversion rather than as final products. Among them, C_26_H_30_O_9,_ which we designate as 10olanoeclepin D (SolD), may act as the direct precursor of the non-methylated form of SolA (C_26_H_28_O_9_, that we coin 10olanoeclepin E, SolE).

### SolA production is affected by soil type

Building further on the importance of the soil microbiome (Figure 2C), we explored the impact of various soils on the conversion of the SolA precursor produced by tomato plants to SolA. Interestingly, tomato plants grown in a Dutch forest soil did not produce any SolA, whereas SolA production was observed in several potting soils (Figure 3A). Plants grown on a mix of forest soil and potting soil also displayed reduced SolA production (Figure 3A) suggesting that the microbes responsible for SolA production are affected. To further explore this, 16S rDNA sequencing was performed on the rhizosphere compartment of plants grown in potting soil and potting soil-forest soil mixtures. A correlation analysis between ASVs abundances with varying SolA levels revealed that both the rhizosphere and root compartment of plants grown in 100% potting soil were enriched in *Massilia* species compared to those grown in forest soil or mixed soil substrates (Figure 3B-C, Figure S5, Table S2).

**Figure 3.**
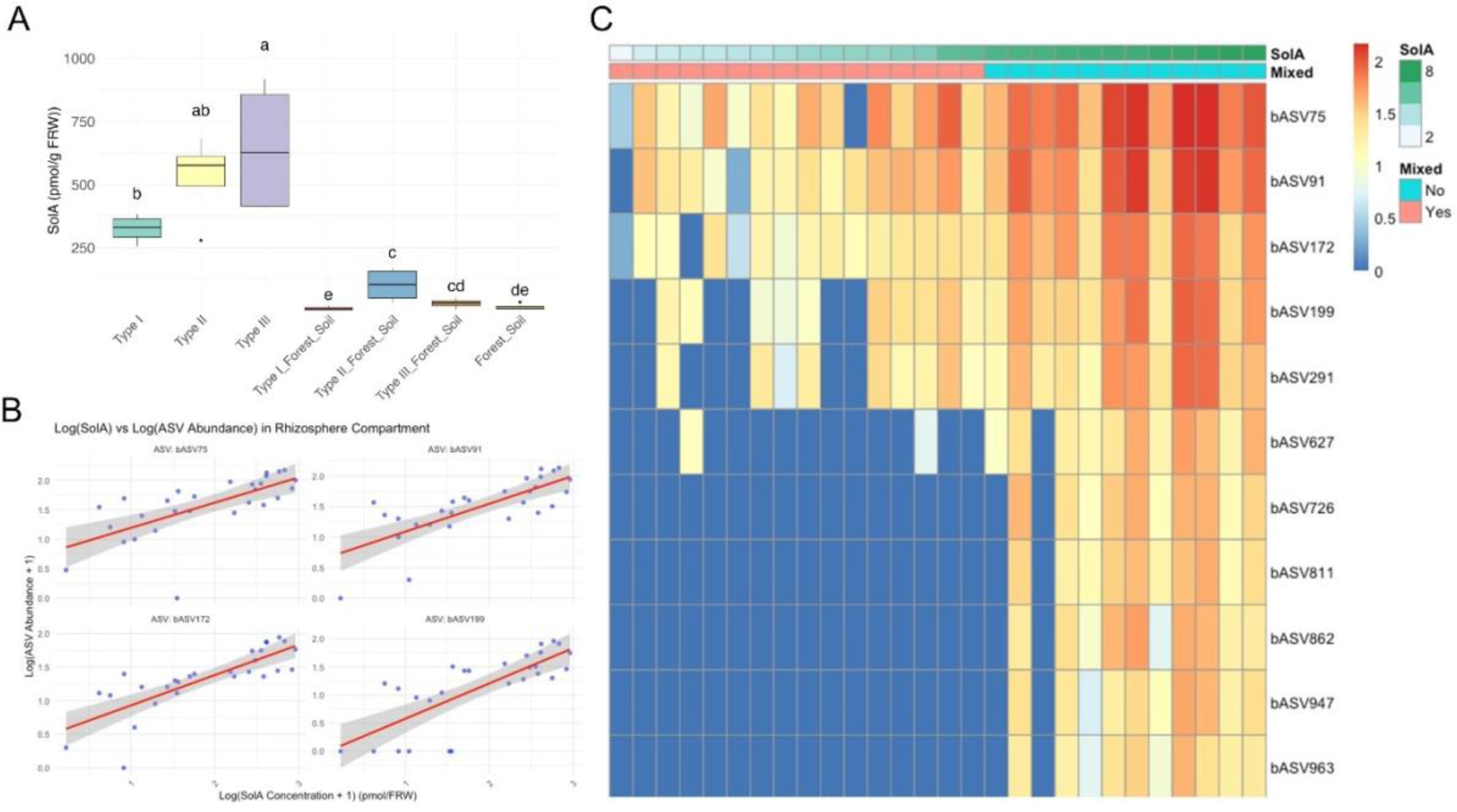
Analysis of SolA production and microbial community composition across different soil types. (A) Boxplot showing SolA concentrations in root exudates of tomato plants grown in different soil types (Type I, II, III, and Forest soil) without and with the addition of 10% forest soil (Type I_Forest, Type II_Forest and Type III_Forest). SolA concentrations are expressed as picomoles per gram of fresh root weight (pmol/g FRW). B) Scatter plots for the top four ASVs assigned to *Massilia spp*. with the highest correlation with the SolA concentration (C) Heatmap of log-transformed abundance of bacterial ASVs assigned to *Massilia spp*. in the rhizosphere, stratified by SolA concentration, forest soil amendment (Mixed) and soil types (soil). The color gradient represents relative ASV abundance, with red indicating higher abundance and blue indicating lower abundance.

To investigate whether *Massilia* spp. contribute to the conversion of the SolA precursor to SolA, *Massilia* strains – isolated from one of the soils that we had shown to stimulate the conversion of the precursor to SolA – were assayed in our conversion assay. However, none of the isolates tested catalyzed the conversion of the precursor to SolA. This result favours the hypothesis that *Massilia* spp. are attracted to the SolA precursor rather than being directly involved in its biosynthesis. To test this hypothesis, we set out to identify SolA (precursor) biosynthetic genes through an RNAseq approach. With these genes in hand we can then analyse the effect of reduced SolA precursor exudation into the tomato rhizosphere on microbiome recruitment, through transient silencing of these genes. The strong effect of N deficiency on SolA biosynthesis (Figure 1B) was used to inform our treatment choice aimed at creating clear differences in SolA production and corresponding gene expression.

### N deficiency induces the expression of SolA biosynthetic gene candidates

To identify SolA biosynthetic gene candidates, we conducted a time-series experiment in the aeroponics system, exposing tomato plants to N deficiency and control conditions, with the seedlings established in soil or rockwool. As expected, N deficiency resulted in a significant increase in SolA content in the exudate of plants grown on soil substrate, while no SolA was detected in the exudate of plants grown on rockwool (Figure S6). Transcriptome analysis on the tomato roots demonstrated a significant effect of both N deficiency and substrate type on gene expression (Figure 4A, Table S3). Differential expression analysis yielded a large number of N-deficiency upregulated genes (Figure 4B, Figure S7, Figure S8), with 346 and 163 genes uniquely upregulated in soil and rockwool, respectively, and 369 upregulated genes shared between the two substrates (Figure 4C). The differences between the substrates may reflect the role of microorganisms in soil, which could influence plant root physiology.

**Figure 4.**
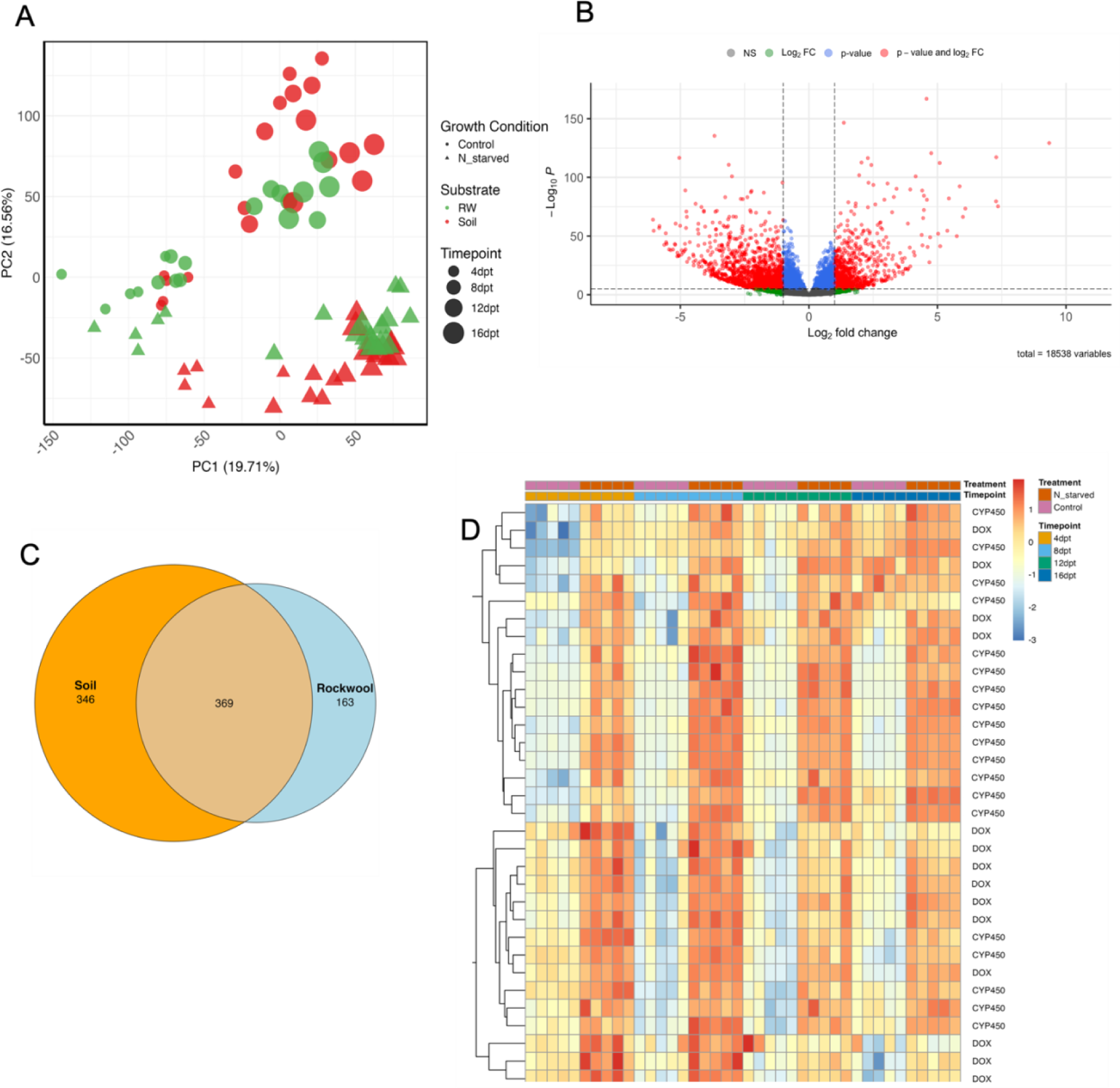
Identification of SolA biosynthetic genes in tomato root transcriptome. A) Principal Component Analysis (PCA) plot representing transcriptional changes in tomato roots under various conditions including substrate, nitrogen availability, and different timepoints. B) Volcano plot illustrating differentially expressed genes (DEGs) with significant changes in expression in soil substrate. C) Venn diagram depicting the number of unique and shared upregulated genes between soil and rockwool substrates under nitrogen starvation. D) Heatmap of SolA candidate genes identified through co-expression analysis with *Solyc06g067870*, *Solyc12g042980*, and *Solyc05g011970* as baits in tomato plants grown on soil. Each gene is assigned to their gene families including 2-oxoglutarate (2OG) and Fe(II)-dependent oxygenase (DOX) and cytochrome P450 (CYP).

To identify candidate genes potentially involved in SolA biosynthesis, we performed co-expression analysis using three previously reported biosynthetic genes (*Solyc06g067870*, *Solyc12g042980*, and *Solyc05g011970*) as baits (Shimizu et al., 2023). Note that Solyc06g067860 and Solyc05g011940 are absent from our dataset thus they were not included as baits. Specifically, we calculated Pearson’s correlation coefficients (PCCs) between all genes and SolA1, SolA3, and SolA4. Genes with a PCC ≥ 0.6 and a *p*-value < 0.05 were considered significantly co-expressed. To continue, we only focused on cytochrome P450 monooxygenases (P450s) and 2-oxoglutarate/Fe(II)-dependent dioxygenases (DOXs) (data not shown). Figure 4D shows 33 *P450s* and *DOX* genes co-expressed with SolA1, SolA3, and SolA4, all of which are upregulated under nitrogen starvation. Moreover, phylogenetic trees were generated for all genes belonging to DOXs and P450s (data not shown). From these, five P450 genes together with *Solyc05g011970* and *Solyc05g011940* were transiently silenced using Virus Induced Gene Silencing (VIGS). Root exudates from transiently silenced and control plants were collected and analyzed for SolA levels. Transient silencing of *Solyc03g114940*, *Solyc05g011970* (*CYP749A20*), and *Solyc05g011940* (*CYP749A19*), which were significantly upregulated under nitrogen starvation (Figure S9), resulted in a significant reduction in the SolA concentration in the root exudate of the silenced plants (Figure S10, S11). The other 4 *P450* genes did not significantly reduce the SolA concentration in the root exudate upon transient silencing, suggesting they may be false positives. This demonstrates that the three cytochrome P450s are involved in the biosynthetic pathway of the SolA precursor. Sequence alignment and phylogenetic analysis indicates that *Solyc03g114940* belongs to the cytochrome P450 subfamily CYP78A, which has been reported in various plant species, primarily within the *Solanaceae* family (Figure S12). The *Solyc03g114940* protein sequence is 100% identical to *CYP78A75* in *Solanum lycopersicum* and shares 95% sequence identity with its ortholog CYP78A75 in *Solanum tuberosum*. Therefore, we designate it as *CYP78A75*.

### Reduced SolA biosynthesis alters the tomato root microbiome composition

The successful reduction in the SolA (precursor) root exudation of VIGS-treated plants allowed us to test the hypothesis that SolA (or its precursor) plays a role in recruiting microbiota. Hereto, we used two genes, *CYP749A20* and *CYP749A19*, which we showed by VIGS to be involved in SolA precursor biosynthesis (Figure 5A, Figure S10, S11) and which had been shown before to be involved in 15olanoeclepin biosynthesis (Shimizu et al., 2023). Using VIGS, these two genes were transiently silenced and the rhizosphere soil of the silenced tomato plants was collected. Microbial DNA was extracted and subjected to 16S rDNA sequencing. While Shannon diversity showed no significant differences between the target gene silenced plants and the GUS control (Figure 5B), silencing SolA biosynthetic genes resulted in a decrease in the relative abundance of several bacterial families, including *Oxalobacteriaceae*, *Flavobacteriaceae*, *Reyranellaceae*, *Xanthobacteraceae*, and *Rhizobiaceae* (Figure 5C and Figure S13). Notably, the abundance of *Massilia* species was reduced in both SolA biosynthetic gene silenced plants (Figure 5D). These results show that SolA or its biosynthetic precursor plays a role in modulating the rhizosphere microbial community, particularly by impacting the abundance of *Massilia* species.

**Figure 5.**
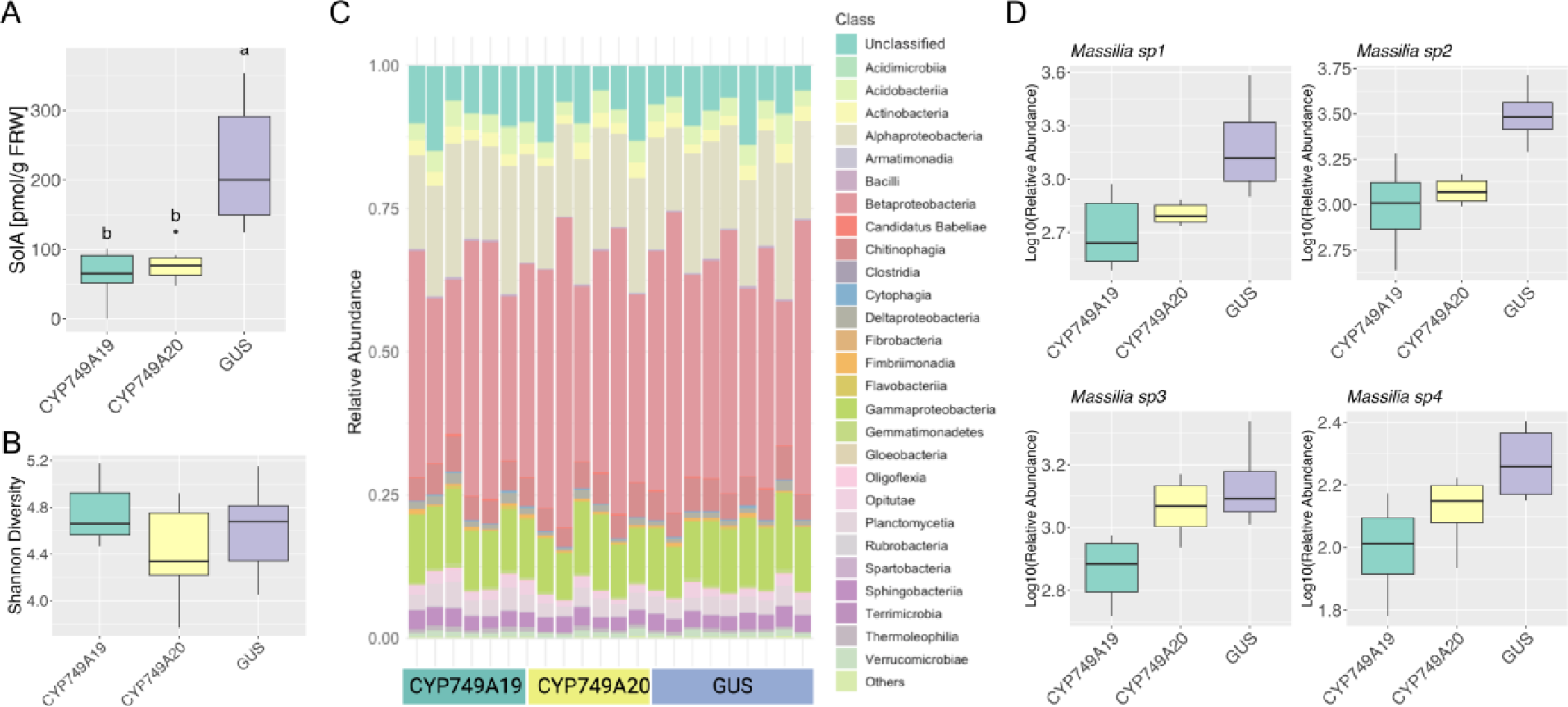
Effects of reduced SolA biosynthesis on the tomato rhizosphere microbiome. (A) SolA concentration in root exudate of tomato plants in which *CYP749A19* and *CYP749A20* were transiently silenced using VIGS, with GUS as a control. SolA concentrations are expressed as picomoles per gram of fresh root weight (pmol/g FRW). (B) Shannon bacterial diversity in the rhizosphere of transiently silenced plants. (C) Stacked bar chart showing the relative abundance of microbial taxa. (D) Boxplots depicting the abundance of selected *Massilia spp*.

### *Massilia sp.* GER05 has plant growth promoting traits

Bacterial profiling of the tomato rhizosphere under nitrogen deficiency revealed marked compositional shifts relative to controls group (Figure S14A-B). Among the enriched taxa, *Massilia* spp. consistently increased in abundance (Figure S14C). Given its positive association with the SolA precursor pathway; enrichment under high SolA and depletion when SolA was low (Figure 3), together with reduced abundance upon silencing *CYP749A19/A20* (Figure 5), we targeted *Massilia* for isolation. A representative isolate was obtained from N-starved rhizospheres and designated GER05. Whole-genome sequencing of GER05 showed that this strain represents *Massilia cellulosiltytica*. The rhizosphere compatibility of GER05 was predicted based on the presence of catabolic gene clusters in the genome that could be involved in catabolism of metabolites in root exudates, such as sugars, amino acids especially tryptophan and polyamines (Figure 6A). Furthermore, the genome of GER05 harbored putative growth promoting genes involved in direct and indirect mechanisms of plant growth promotion such as those responsible for colonization and competition for resources in the rhizosphere (Figure 6B). The genome also reveals genes associated with the production of phytohormones, including indole-3-acetic acid (IAA). This was confirmed by experimental data (Figure 6C), further supporting a role in promoting plant growth. Additionally, the bacterial genome harbored nitrogen acquisition and phosphate solubilization associated genes (Figure S15), which are critical for enhancing plant resilience and adaptation to nutrient-limited conditions. To assess whether these genomic traits translate into plant growth benefits, a greenhouse experiment was conducted to evaluate the impact of GER05 on tomato growth under nitrogen deficiency. Uninoculated plants displayed poor growth in the substrate with limited N, including poor root growth and chlorotic leaves (Figure 6D). Inoculation with GER05 resulted in improved plant growth compared with the control, with enhanced root and shoot growth as well as chlorophyll content (Figure 6D).

**Figure 6.**
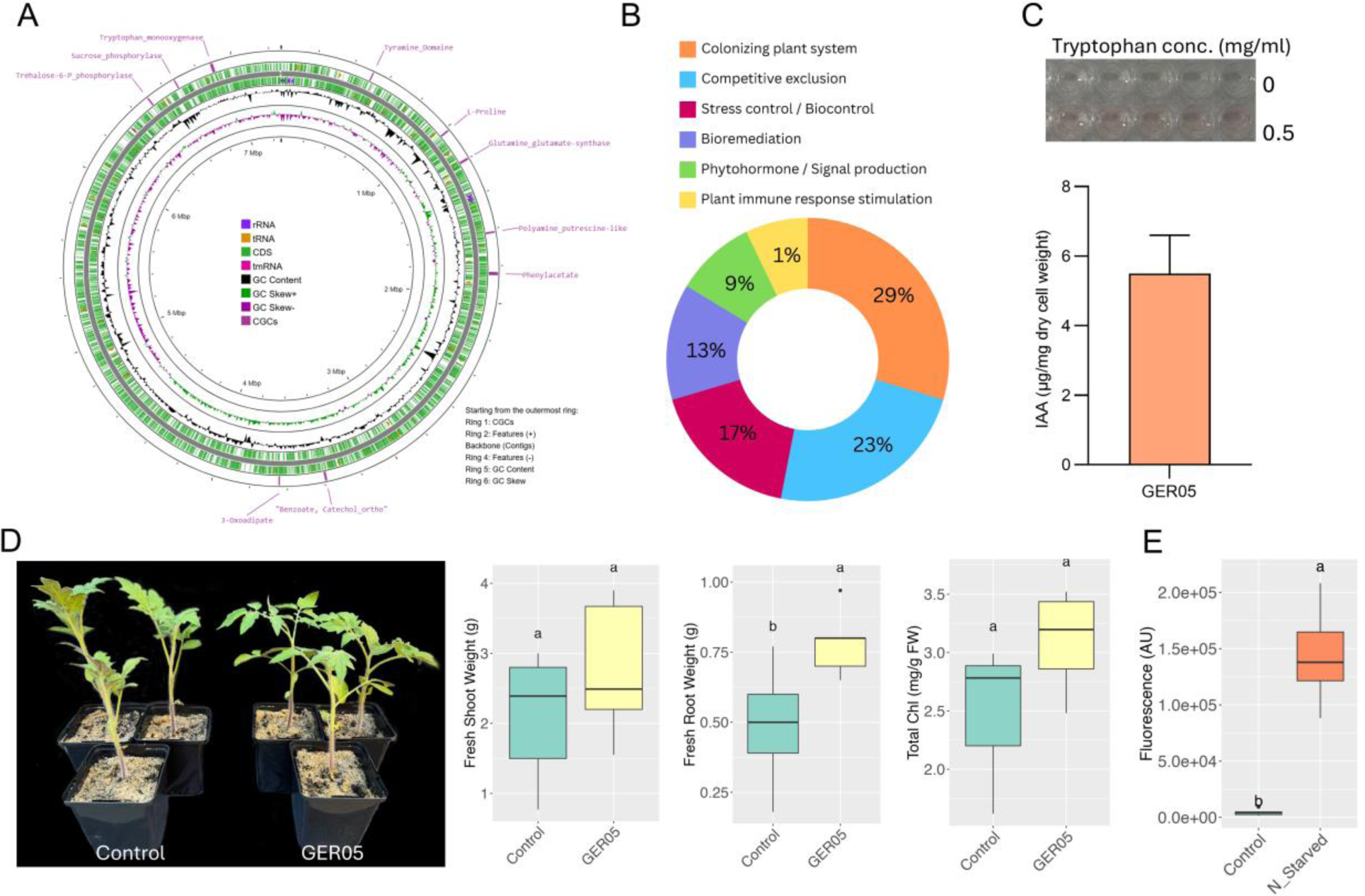
Genomic features and plant growth promoting effect of *Massilia cellulosiltytica* GER05. (A) Genome map of *Massilia* GER05 showing RhizoSMASH predicted catabolic gene clusters (CGCs, outermost ring) and other features. (B) Prediction of overall plant growth promoting traits (PGPT_pred) in the genome of GER05 based on PlaBAse database. (C) Validation of indole acetic acid production *in vitro* in the presence of tryptophan as precursor. (D) Effect of GER05 on the growth of tomato under N-deficiency after two weeks of starvation. € Chemotaxis assay showing the attraction of GER05 to root exudates of tomato plants grown on rockwool N starved and control conditions.

To further strengthen our mechanistic understanding, we evaluated whether IAA alone could account for the growth-promoting effects of *Massilia* sp. GER05. Exogenous application of IAA (0.5 and 5 mg/L) to N-starved tomato plants increased the root-to-shoot ratio, indicating enhanced biomass allocation to roots. GER05 treatment similarly increased the root-to-shoot ratio, consistent with its IAA production, but additionally promoted greater total biomass compared to IAA treatments alone (Figure S16A-B). Moreover, GER05-treated plants displayed a trend toward higher shoot nitrogen content relative to both N-starved and IAA-treated plants (Figure S16C). This suggest that GER05 may improve nitrogen availability, potentially by stimulating plant nitrogen uptake systems or by enhancing mineralization processes that release organic nitrogen into plant-available forms. Although these differences were not statistically significant, the pattern aligns with earlier chlorophyll and leaf greenness measurements (Figure 6D), supporting the view that GER05 promotes plant growth under nitrogen-limited conditions through a combination of IAA-mediated root modulation and enhanced nitrogen availability.

Finally, a chemotaxis assay with the isolated GER05 and root exudates from tomato plants grown on rockwool under N starved and control conditions showed that GER05 is strongly attracted towards the N starved root exudates (Figure 6E). On the other hand, the chemotaxis assay conducted with SolA and GER05 revealed no observable attraction of the GER05 strain towards SolA (Figure S17). These results suggest that the SolA precursor, rather than SolA itself, is likely responsible for the attraction of *Massilia*.

## Discussion

Triterpenoid eclepins, such as SolA and glycinoeclepins, were discovered as hatching stimulants for parasitic PCN and soybean cyst nematode (SCN), respectively (Masamune et al., 1982; Fukuzawa et al., 1985; Mulder et al., 1996). In the present study, we postulate that these eclepins must have a so far unknown beneficial role for plants. We show that the biosynthesis of SolA is induced by N deficiency through the upregulation of biosynthetic gene expression, and we demonstrate that SolA is jointly produced by tomato, which produces a SolA precursor, and soil microbiota, which convert this precursor to SolA. Transient silencing of SolA biosynthetic genes resulted in a strong decrease in SolA exudation by tomato, and a concomitant, reduced recruitment of *Massilia* spp. to the tomato rhizosphere. An isolate of *Massilia cellulosiltytica*, obtained from the rhizosphere of tomato grown under N deficiency, displays growth promoting activity in tomato exposed to N deficiency.

Our finding that SolA production involves both plant enzymes and soil microbiota is striking. Tomato plants grown in (semi-)sterile substrates (rockwool and gamma-irradiated soil) did not produce SolA (Figure 2A). However, when the root exudates from these plants, which initially lacked SolA, were incubated with soil, SolA could be detected. Mulder *et al*. (1996) were the first to report the isolation of SolA, extracting it from the root exudate of 700 potato plants grown in a hydroponic system (Mulder et al., 1996). Later, Guerrieri *et al* developed an advanced analytical method to detect this molecule in the root exudate of a single plant growing in soil substrate (Guerrieri et al., 2021). Since SolA was detected in both studies, both must have conducted their research under non-sterile conditions. Here we present evidence that the plant itself is unable to produce SolA. In other words, the plant exudes a SolA precursor into the rhizosphere where soil microbes convert it into SolA. Recently, Shimizu et al identified 19olanoeclepin B (SolB), a compound structurally related to SolA, as a novel hatching factor isolated from potato hydroponic culture and tomato hairy roots (Shimizu et al., 2023). When this SolB was incubated with soil, it was converted to SolA. Similarly, here we demonstrate that root exudate of tomato plants grown on rockwool does not contain SolA, but when this root exudate is incubated with soil, SolA production occurs. However, unlike the findings of Shimizu et al., SolB was virtually absent in our tomato root exudates.

SPE column-based fractionation showed that the SolA precursor already contains the carboxy group. Moreover, untargeted metabolomics of the optimized conversion assay demonstrated that when root exudates from tomato plants grown in rockwool were incubated with non-sterile soil, SolB appeared. This was also the case for C_26_H_28_O_9_, SolC. The discrepancy between our findings and those of Shimizu et al. is likely due to differences in experimental systems. While Shimizu et al. used tomato hairy roots, we analyzed whole tomato plants. Hairy roots often exhibit distinct metabolic profiles compared to intact plants, and their generation involves *Agrobacterium rhizogenes*, which may influence metabolite secretion or even play a role in SolB production. In addition, we detected small amounts of C_26_H_30_O_9_ (SolD) when root exudates from rockwool-grown plants were incubated with sterile soil (Table S1). A slight but non-significant increase was observed in non-sterile soil, suggesting that the plant might secrete SolD, which may serve as a direct precursor for microbial conversion into the non-methylated form of SolA (SolC, C_26_H_28_O_9_). We hypothesize several pathways for SolA biosynthesis that use SolB, SolC, acetylated SolB, and/or C27H34O7 as intermediates, with participation from one or more microbial species. We are currently isolating both the individual intermediates and the microbes responsible for their conversion.

In a next step, we confirmed our hypothesis that SolA, or as it now turns out its precursor, must have a beneficial role for the plant. First, using metabarcoding analysis we revealed that tomato plants grown on potting soil, on which they produce high levels of SolA, have a rhizosphere enriched with *Massilia* spp. However, when grown on forest soil or a mixture of potting compost and forest soil, SolA production was virtually negated, while there was a corresponding decrease in *Massilia* spp. abundance. We first tested whether this implied that *Massilia* spp. is the micro-organism that converts the SolA precursor to SolA, but this was not the case, making it likely that the SolA precursor is a recruitment signal for *Massilia* spp.

This was further confirmed by disrupting the exudation of SolA precursor into the rhizosphere, through VIGS. Transcriptome analysis of tomato roots under nitrogen starvation conditions identified candidate SolA biosynthetic genes. Silencing these genes, including *CYP749A20* and *CYP749A19*, not only reduced SolA accumulation but also altered the root microbiome, specifically reducing the abundance of *Massilia* spp., further supporting the role of SolA precursor in recruiting *Massilia* spp. Isolation of enriched bacteria from the rhizosphere of tomato plants grown under nitrogen deficiency in soil resulted in identifying *Massilia* sp. GER05. Inoculation of tomato plants with GER05 under nitrogen-deficient conditions enhanced plant growth and increased chlorophyll content (Figure 6). Moreover, whole-genome sequencing confirmed the presence of IAA biosynthesis genes, supporting the experimental evidence that GER05 can produce IAA. The *Massilia* sp. GER05 genome also harbors genes involved in nitrogen acquisition, nitrogen mineralization and phosphate solubilization, which may contribute to improved nitrogen uptake under nutrient-deficient conditions. In line with this, complementary experiments revealed that exogenous IAA treatments partially mimicked the effects of GER05 on root-to-shoot allocation, but GER05 inoculation promoted greater overall biomass, indicating that its effects cannot be explained by IAA production alone. Moreover, tomato plants treated with GER05 tended to accumulate more shoot nitrogen compared with IAA-treated or uninoculated controls. These findings suggest that GER05 promotes plant growth under nitrogen limitation through multiple mechanisms, with IAA-mediated modulation of root architecture complemented by enhanced nitrogen availability in the rhizosphere. Moreover, Han et al. (2024) recently demonstrated that *Massilia*-treated soybeans upregulate auxin-related genes in both roots and leaves, reinforcing our conclusion that exogenous IAA application alone does not fully reproduce the effects observed in *Massilia* (GER05)-treated plants (Han et al., 2024). Interestingly, the SolA concentration in tomato root exudates increases under nitrogen- and, to a lesser extent, phosphate-deficiency, which consequently resulted in more PCN hatching. A chemotaxis assay confirmed that GER05 is attracted towards root exudates from nitrogen-starved tomato plants grown in rockwool, but it did not show a response to SolA. These findings indicate the hypothesis that a SolA precursor, rather than SolA itself, is probably responsible for recruiting beneficial microbes including *Massilia* spp. Indeed, transcriptomic analyses in tomato and Arabidopsis have shown that exogenous SolA treatment downregulates plant immune and stress responses while promoting root growth and altering root architecture, particularly under nutrient deficiency (Vlaar et al., 2022). These findings suggest that SolA not only functions as a nematode hatching factor but also acts as a signaling molecule shaping plant–microbe interactions. To disentangle the direct rhizosphere signaling roles of SolA and its precursor from indirect physiological effects, future experiments such as blocking SolA transport or applying exogenous SolA and its precursor will be essential. *Massilia* spp. have been shown to have plant growth-promoting effects, enhancing nitrogen uptake and supporting maize growth under nitrogen-limited conditions (Yu et al., 2021; Wang et al., 2024). Moreover, Krishnamoorthy et al isolated *Massilia* sp. RK4 from the spores of *Rhizophagus intraradices*, which plays a crucial role in enhancing maize salt stress tolerance. Co-inoculation with *R. intraradices* and *Massilia* sp. RK4 improved AMF root colonization, nutrient accumulation, and overall plant growth under saline conditions (Krishnamoorthy et al., 2016). Here, we demonstrate that *Massilia* sp. GER05 is attracted to a precursor of the triterpenoid SolA, and that disruption of the corresponding biosynthetic genes in plants impairs *Massilia* spp. recruitment. This work provides a foundation for further investigations into the identity of the precursor involved and may facilitate the discovery of similar attractants for *Massilia* spp. in other crop species. The growth promoting activity of *Massilia* spp. under conditions of reduced N availability opens up possibilities for reducing the use of vast amounts of nitrogen fertilizer in agriculture, potentially reducing the negative impact of agriculture on the environment, and making it more sustainable.

The interaction between host organisms and microbes In the production of joint bioactive compounds represents an emerging area of study. Our findings indicate that tomato plants rely on soil microbes to complete the biosynthesis of SolA from a plant-produced precursor—a clear example of cooperative metabolism (Figure 7). Under nitrogen deficiency, tomato plants produce higher levels of the SolA precursor, effectively sending a “cry for help” message that recruits beneficial *Massilia* spp. that help to improve nitrogen acquisition and possibly other microbes. One or more of these microbes convert the SolA precursor into SolA, which is then exploited by cyst nematodes. However, we propose that SolA also benefits the microbes themselves by lowering host plant immunity and facilitating colonization, as hinted by Vlaar et al. (2022), who demonstrated that SolA treatment reduces the expression of immunity-related genes, especially under nitrogen deficiency (Vlaar et al., 2022b).

**Figure 7.**
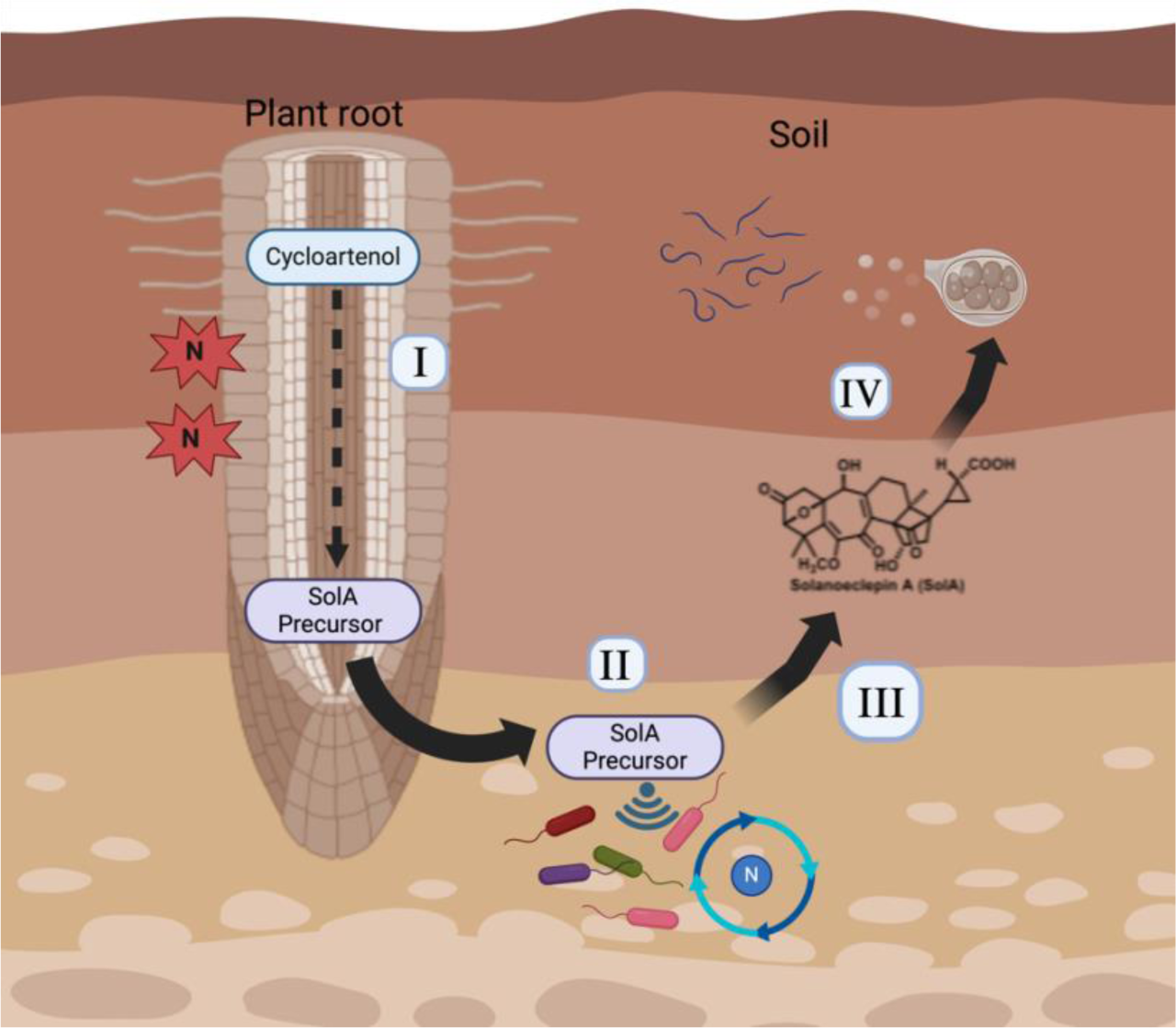
Conceptual framework illustrating the role of nitrogen deficiency and microbiota in Solanoeclepin A (SolA) biosynthesis and plant-microbe interactions. (I) Under nitrogen (N) deficiency, tomato roots exude a SolA precursor, which is derived from cycloartenol through an upregulated biosynthetic pathway. (II) The exuded SolA precursor attracts beneficial rhizosphere microbes, including *Massilia* spp. (III) The resulting attracted microbiota improve plant growth under nitrogen deficiency. (IV) Microbiota further converts the SolA precursor into SolA, which is hijacked as a host presence cue by potato cyst nematodes (PCNs). This figure was created in BioRender (https://biorender.com/).

An interesting parallel exists in animal systems, where gut microbes transform steroidal primary bile acids – such as cholic and chenodeoxycholic acid – into secondary bile acids like deoxycholic and lithocholic acid. These microbial modifications significantly alter the biological activity of bile acids, impacting lipid metabolism, shaping gut microbiome composition, and modulating immune responses via nuclear receptors (Dawson and Karpen, 2015; Guzior and Quinn, 2021; Lee et al., 2022). This type of host-microbe co-metabolism suggests a case of convergent evolution, where both plants and animals leverage microbial transformations to fine-tune bioactive signaling compounds for improved ecological interactions. The dual functionality of SolA inducing hatching of PCN and its precursor being a recruitment signal for *Massilia* spp. is reminiscent of the biological activity of the strigolactones, which are recruitment signals for beneficial arbuscular mycorrhizal fungi to enhance nutrient uptake, while being abused as germination stimulants by root parasitic Orobanchaceae (Akiyama and Hayashi, 2006; Bouwmeester et al., 2021). Also, the biosynthesis of strigolactones is enhanced by nutrient deficiency, particularly phosphate and, to a lesser extent, nitrogen (Al-Babili and Bouwmeester, 2015). There are other rhizosphere metabolites such as benzoxazinoids and glycoalkaloids that are also subject to microbial conversion (Kudjordjie et al., 2019). However, not all microbial transformations necessarily imply a signaling function. The latter metabolites are produced in much higher concentrations, suggesting they may primarily function as antimicrobial defense compounds rather than precise microbial recruitment signals (Bouwmeester et al., 2025).

In conclusion, this study advances our understanding of the biological relevance of SolA and demonstrates its dual role in the rhizosphere, especially under nitrogen-limited conditions. A SolA precursor is a signal that attracts beneficial microbes, most likely *Massilia* spp., which support nutrient acquisition, while SolA itself functions as a hatching stimulant for PCN. We anticipate that the biological relevance of these findings extends to the glycinoeclepins, which share a strong structural similarity and stimulate hatching in soybean cyst nematode. This shared functionality suggests that SolA and glycinoeclepins may not only have related biosynthetic pathways but also analogous roles in shaping the rhizosphere interaction of plants with microbiota across the plant kingdom. Together, these findings position SolA – and perhaps the eclepins in general – as a multi-functional regulator of the interaction of plants with other organisms in their rhizosphere. In future work we will further resolve the SolA biosynthetic pathway, identify the microbes converting SolA precursor to SolA and explore the possibility to harness the eclepins for enhancing crop resilience and managing pest pressure in sustainable agricultural systems.

## Material and methods

### Tomato growth conditions and nitrogen and phosphorus treatments

Surface-sterilized tomato (*Solanum lycopersicum* cv. Money maker) seeds were initially grown in an aeroponic system using complete Hoagland solution for two weeks to establish seedlings. Following this initial growth period, different nitrogen treatments were applied by replacing the existing Hoagland solution with a modified solution based on the experimental needs, while maintaining a control group (with a final nitrogen concentration of 11.2 mM/L) with complete Hoagland solution across all experiments.

For the nitrogen-experiments, NH₄NO₃ was removed from the Hoagland solution while another group received double amount of nitrogen (final concentration of 11.2 mM/L). Root exudates from these groups at multiple time points for SolA quantification. Moreover, to evaluate the effects of phosphorus deficiency, K_2_HPO_4_ was removed from the Hoagland solution. To compensate K deficiency, similar amount was replaced by adding KCl.

In the nitrogen repletion experiment, two-week-old plants were first subjected to nitrogen starvation by replacing the complete Hoagland solution with an NH₄NO₃-free version for two weeks. After this starvation period, one subset of nitrogen-starved plants was returned to complete Hoagland solution to allow for nitrogen recovery, while another subset remained nitrogen-starved. Root exudates were collected from the remaining solution at each replacement and then at 5 and 10 days following nitrogen repletion to assess the effects of nitrogen recovery.

To examine the effects of separate nitrogen sources, tomato plants were also subjected to different nitrogen conditions: nitrate-free, ammonia-free, and completely nitrogen-free. In the nitrate-free condition, NH₄NO₃ in the Hoagland solution was replaced with KCl to maintain ionic balance. For the ammonia-free condition, NH₄NO₃ was replaced with Ca(NO₃)₂ to provide nitrate without ammonia. In the completely nitrogen-free condition, NH₄NO₃ was entirely removed from the Hoagland recipe. Root exudates were collected from each treatment group, including the control, after one and two weeks of nitrogen deprivation.

In all conditions described above, 9 mL of remaining Hoagland solution was used for SolA analysis. Moreover, fresh root tissues and the volume of remaining Hoagland solution from each boucket were considered for SolA quantification.

### SolA extraction and UPLC-MS/MS analysis

SolA analysis was conducted as described by (Guerrieri et al., 2021). Briefly, the pH of filtered root exudates was adjusted to a range of 6.9-7.2. Solid-phase extraction was performed on root exudate solutions with a polymeric mixed mode anion exchanger (SPE Oasis® MAX; 3 cc/60 mg, Waters, Milford, MA, USA). The cartridges were preconditioned with 3 mL of 100% methanol, 3 mL of a 5% NH_4_OH/H_2_O (v/v) solution (pH 11) and 3 mL of Milli-Q water. Then 5 mL of the pH adjusted root exudate was loaded onto the preconditioned cartridges. The cartridges were washed with 3 mL of Milli-Q water and 3 mL of 100% methanol. The analytes were eluted with 3 mL of 0.2 M Formic acid in methanol, and the eluate was subsequently evaporated in a Speed Vacuum Concentrator (ScanSpeed 40, LaboGene, Denmark). The dried extract was reconstituted in 120 μL 20% methanol, then filtered through micro-spin nylon filter (0.2 μm pore size, Thermo, Waltham, MA, USA).

The analysis of SolA was performed using a Waters Acquity ultra-high pressure liquid chromatography (UPLC) I-Class System (Waters, Milford, MA, USA) equipped with a binary solvent manager and sample manager, coupled to a Xevo® TQ-XS tandem quadrupole mass spectrometer (MS/MS, Waters MS Technologies, Manchester, UK) with electrospray (ESI) ionization interface. A 5 µL aliquot of the extracted sample was injected onto the UPLC column (Acquity UPLC Ethylene Bridged Hybrid (BEH) C18 column, 2.1 x 100 mm, 1.7 μm particle size, Waters, Milford, MA, USA), kept at a constant temperature of 40°C. The analyte was eluted at a flow rate of 0.3 mL/min using a 9 min linear gradient of 15 mM formic acid/water (A) and 15 mM formic acid/acetonitrile (B) with the following elution profile: 0-1 min (5% B), 6 min (50% B), 8 min (80% B), 9 min (95% B). At the end of the gradient, the column was washed with 95% solvent B for 1 min and finally equilibrated to initial conditions for 2 min. The analyte was introduced into the ESI ion source of the mass spectrometer and SolA was analyzed in positive mode as [M+H]^+^, using diagnostic and confirming precursor-to-product transitions 499>83, >399, >315, >453 with optimized collision energy 30, 25, 25, and 20 eV, respectively, and collision gas (argon) flow of 0.15 mL/min. The instrument control, MS data acquisition, and processing were carried out by the MassLynx software, version 4.2 (Waters). After determining SolA concentration in the extracted sample, the amount of SolA was quantified per gram of roots.

### Bioconversion assay and fractionation of root exudates using ion exchange cartridges

Tomato plants were grown on rockwool substrate for four weeks under nitrogen starvation in an aeroponic system as described above. After the growth period, 50 mL of the remaining root exudate solution was collected and concentrated fivefold using a freezedryer. A 5 mL aliquot of the concentrated solution was then incubated with both autoclaved (sterilized) and unsterilized MiCrop soil samples at 25°C for 48 hours. Following incubation, the liquid phase was carefully extracted from each tube, filtered, and subjected to solid-phase extraction (SPE) for SolA isolation and quantification. The presence of SolA was subsequently analyzed using UPLC-MS/MS.

To fractionate the root exudates, Oasis® MAX (3 cc/60 mg) cartridges were preconditioned by sequentially washing with 6 mL of 100% methanol, 6 mL of a 5% NH₄OH/H₂O (v/v) solution (pH 11), and 6 mL of Milli-Q water to ensure optimal binding conditions. 20 mL of pH-adjusted root exudate was then passed through the preconditioned cartridge. Elution was performed stepwise using Milli-Q water, 100% methanol, and 0.2 M formic acid in methanol, with the flowthrough collected after each step. The water fraction was dried using a freeze dryer, while the methanol and formic acid in methanol fractions were evaporated using a SpeedVac concentrator.

The resulting pellets from each fraction were dissolved In 100 µL of pure ethanol, followed by the addition of 4 mL of sterilized water. These prepared solutions were incubated with unsterilized soil for 48 hours to facilitate potential interactions or conversions of precursor compounds by soil microorganisms. After the incubation period, SolA was extracted from the soil using SPE and analyzed using UPLC-MS/MS to detect the presence of SolA. Crude root exudates were also incubated with sterilized soil as negative control.

MiCrop soil was collected from an organically managed agricultural field in Nergena, Bennekom (coordinates: 51.996250, 5.659375). It was collected in 2014 by excavating the 80 cm top layer of the field that was subsequently stored outdoors and left unmanaged, thereby ensuring that a natural soil community was sustained through the presence of wild plants. Before use in experiments, batches of the soil were air-dried and sieved through a 5 mm mesh to remove bigger particles and roots and stored in the dark until use.

### Metabolite profiling and SolA intermediate identification

To investigate the intermediates involved in SolA biosynthesis, root exudates were collected from tomato plants grown on rockwool under N starved conditions and incubated with sterilized and unsterilized soils, as described in bioconversion assay. Samples were loaded into Oasis MAX column and eluted with 0.2M FA in methanol, and after concentrating with SpeedVacs, the resulting pellets were dissolved in 20% methanol (methanol: water) and analyzed using quadrupole time-of-flight mass spectrometer (QTOF) equipped with a dual-stage trapped ion mobility separation cell (timsTOF pro Bruker Daltonics Inc., Billerica, MA, USA). A 20 μL sample injection and liquid chromatography (LC) separation was conducted using the Ultimate RS UPLC system (Thermo Scientific, Germeringen, Germany). The system employed an Acquity UPLC CSH C18 column (130 Å, 1.7 µm, 2.1 mm × 100 mm) coupled with a VanGuard pre-column (2.1 mm × 5 mm) of identical material. A gradient elution was performed, transitioning from 1% to 99% acetonitrile over 18 minutes. Compounds were ionized in both positive and negative ion modes using an Apollo II ion funnel ESI source (Bruker Daltonics Inc.). The data generated were processed and analyzed with DataAnalysis version 4.3 and MetaboScape® 5.0 software, both from Bruker Daltonics.

Metabolic features were filtered based on molecular mass and formula annotation, retaining those within the range of 426-600 Da and containing 26-30 carbon atoms. Features with formulas exclusively composed of C, H, and O were prioritized for subsequent analysis. Key metabolites, including the final product SolA (C₂₇H₃₀O₉), its direct precursor (C₂₇H₃₂O₉) and its acetylated form (C_29_H_34_O_10_), and a potential precursor (C₂₆H₂₈O₉), were annotated in the dataset, despite lacking MS2 spectra, due to their low abundance. These metabolites were included because of their relevance to putative biosynthetic pathways and support from findings in previous publications (Guerrieri et al., 2021; Shimizu et al., 2023).

### RNA sequencing and selection of SolA biosynthetic gene candidates

Tomato seeds were surface-sterilized and grown on a complete Hoagland solution in an aeroponic system with rockwool and unsterilized potting soil as substrates. After 10 days, NH_4_NO_3_ was excluded from the Hoagland solution of N starved groups. Time-series sampling was conducted at 4, 8, 12, and 16 days post-treatment (dpt) from the hanging roots, which were promptly snap-frozen in liquid nitrogen. The collected roots were ground using a mortar and pestle which were held frozen with liquid nitrogen. RNA was extracted from 100 mg of frozen root tissue using Trizol reagent (Sigma), followed by chloroform purification and an RNAeasy Plant Mini Kit (Qiagen). Potential genomic contamination was digested using Dnase (Rnase-Free Dnase Set, Qiagen). The quality and concentration of the extracted RNA were measured using a NanoDrop spectrophotometer. Only high-quality RNA samples, with an OD 260/280 ratio greater than 1.8 and an OD 260/230 ratio exceeding 2.0, were selected for sequencing.

Data preprocessing was performed using a Snakemake (v5.2) pipeline (Köster and Rahmann, 2012). Read quality was assessed and trimmed with fastp (v0.19.5) (Chen et al., 2018) to remove low-quality sequences and adapter contamination. The processed reads were then aligned to the *Solanum lycopersicum* SL3.0 reference genome from the Solanaceae Genomics Project (2018) using STAR (v2.7.6a) (Dobin et al., 2013), ensuring high-accuracy mapping. Aligned reads were assigned to transcript IDs using featureCounts (v1.6) from the Subread package (Liao et al., 2013). Downstream analyses were conducted In R (v4.1). To ensure data quality, raw counts were filtered to retain only transcripts with non-zero expression in all replicates across at least four experimental conditions. Counts were normalized for sequencing depth and transformed using DESeq2 (Love et al., 2014).

Root transcriptome of tomato plants under nitrogen deficient and control conditions were compared to identify differentially expressed genes (DEGs) using DESeq2 package in R. For this end, DEGs in nitrogen-starved vs control in each experimental groups including substrates and timepoints were analyzed separately. DEGs with a log2fold change greater than 2 and an adjusted *p*-value less than 0.001 were selected. Considering the current literature stating that SolA is most likely stems from cycloartenol and involves multiple oxidation and hydroxylation steps (Holden, 2015; Shimizu et al., 2023), the list of selected DE genes were further refined by focusing on cytochrome P450s and 2-oxoglutarate dependent oxygenases.

After validation of *Solyc05g011970* (*CYP749A20*), *Solyc05g011940 (CYP749A19*), a co-expression analysis was performed with MASCARA (White et al., 2024) using one of these Cytochrome-P450s (*Solyc05g011970*) with 2-oxoglutarate (2OG) and Fe(II)-dependent oxygenase (DOX) (i.e. *Solyc06g067870*, *Solyc12g042980*) identified by Shimizu et al., (2023) as baits.

### Virus Induced Gene Silencing

First-strand cDNA synthesis was performed with RevertAid H Minus Reverse Transcriptase (Thermo Fischer Scientific) using 800 μg of the extracted RNA and oligo-dT. By using SNG VIGS TOOL (https://vigs.solgenomics.net/), the best target region to interfere expression of each gene was identified and PCR amplified using specific primers that included attB sequences at their 5’ ends (Supplementary Table S4). The amplicons were cleaned up using GeneJET PCR Purification Kit (Thermo Fisher Scientific, Lithuania) and cloned into pTRV2b vector using gateway cloning. The inserted fragment was validated via PCR amplification and sanger sequencing. pTRV2b construct containing a fragment of a gene of interest was then introduced into *Agrobacterium tumefaciens* strain GV3101 via heat-shock transformation approach.

*A. tumefaciens* strains harboring pTRV1 and pTRV2b constructs were cultured in liquid LB medium supplemented with 20 µM acetosyrigone, 20 mg/l rifampicin and 50 mg/l kanamycin for 2 days in a shaker incubator at 28°C. Bacterial cells were collected by 3000 g centrifugation for 10 min and then resuspended in infiltration solution (10 mM MES pH 5.6, 0.5% MS basal medium and 20 g/L sucrose and 200 μM acetosyringone) to an OD of 1. Each bacterial strain containing pTRV2b carrying one of the targeted gene was mixed 1:1 (v/v) with *A. tumefaciens* containing pTRV1 and incubated at 25°C and darkness condition for 2 h before infiltration. The final suspension was infiltrated into 9-day-old tomato cotyledons using a needless syringe. In this experiment, we also used pTRV2-SlPDS containing a fragment of the *phytoene desaturase* gene from *S. lycopersicum* as visual marker of VIGS efficiency based on photobleaching. Moreover, the β-glucuronidase (GUS) gene was included as a non-target control to distinguish virus effects from target-specific silencing. Infiltrated plants were then grown under an 8/16 h dark/light regime and watered regularly. After four weeks of infiltration, root exudates and tissues were collected for metabolite and transcriptional analysis.

### Chemotaxis assay of *Massilia* sp. GER05 towards SolA and root exudates

The chemotactic assay of GER05 was performed using different concentrations of pure SolA (100 nM and 1 µM). Additionally, crude root exudates from tomato plants grown on rockwool under N starvation (N_starved) and N sufficient (Control) conditions were also used for chemotaxis assay knowing that these exudates do not possesses SolA but its precursor. UPLC-MS/MS confirmed that neither exudates contained SolA. For chemotaxis, overnight cultured GER05 in R2A broth were pelleted at 3500 × *g*, washed, and resuspended in R2A broth (0.1×) to an optical density at 600 nm (OD_600_) of 1. The chemotaxis assay using bait and trap method (Jain et al., 2023) was performed at ambient temperature for 3 h with tips containing SolA or root exudates (200 µL) inserted into a deep 96-well plate containing 1 mL bacterial suspension. Following incubation, tips containing SolA or root exudates were collected and transferred into a 96-well black plate. To quantify chemotactic response of bacteria, cells were stained with 1× BactoView™ Live green fluorescent stain (Biotium, USA) and incubated at 30°C for 30 minutes in dark. Fluorescence intensity was measured using a Synergy H1 microplate reader (BioTek) with an excitation/emission of 485/520 nm. Sample control for SolA contained the same amount of solvent (ethanol) that was used to prepare SolA dilutions and chemotaxis for sample control was also performed along with. Sample blank for SolA and root exudates without chemotaxis were also considered for fluorescence measurement and subtracted from sample fluorescence before final calculations.

## Data availability

The datasets will be made publicly available upon publication in a peer-reviewed journal.

## Authors contributions

DA, LD, and HJB conceptualized and designed the research. DA, AG, and RJ conducted the experiments, and FW performed RNA-seq preprocessing and co-expression analysis. YY and JK assisted with the VIGS and aeroponics setups. GK carried out the untargeted metabolomics. DA, LD, and HJB drafted the manuscript, with all authors contributing to revisions and approving the final version.

## Acknowledgment

The authors acknowledge funding from the European Research Council (ERC) Advanced grant CHEMCOMRHIZO 670211 (HJB), the Dutch Research Council (NWO/OCW) Gravitation program Harnessing the second genome of plants (MiCRop) 024.004.014 (HJB, LD), Marie Curie fellowship NEMHATCH 793795 (LD), the Dutch Research Council (NWO-TTW) Chemical communication between potato and cyst nematodes 16873 (HJB, LD), the Dutch Research Council (NWO-ENW) Vidi grant DECODE VI.Vidi.223.088 (LD), the Data Science Centre of the University of Amsterdam (FW). We thank Professor David R. Nelson (The University of Tennessee Health Science Center) for naming the P450 CYP78A75. We are also grateful to Dr. Aska Goverse (Wageningen University) for her support with the hatching assay.

## Supplementary Information

**Document S1** includes Figures S1-S17, Tables S1-S4, and extended Materials and Methods.

