## Supplementary material for "A Solanoeclepin A precursor functions as a new rhizosphere signaling molecule recruiting growth-promoting microbes under nitrogen deficiency": Document S1

### Supplementary Information

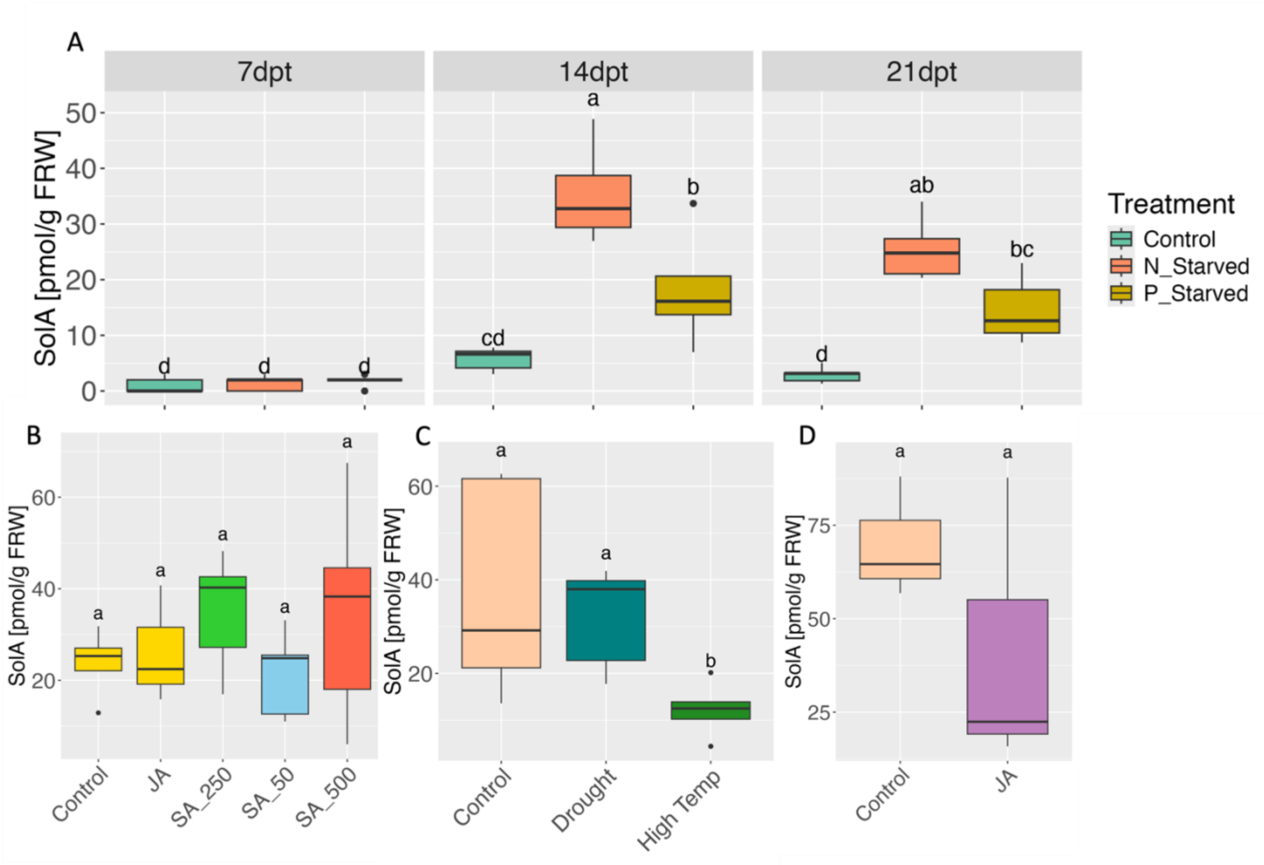


**Figure S1.** Effects of various stresses on SolA production. A) SolA accumulation in root exudates of tomato plants treated with phosphorus and nitrogen deficiency after 7, 14 and 21 days post treatments (dpt) ; B-D) SolA accumulation in root exudates of potato plants under drought, high temperature, Salicylic acid (SA) and jasmonic acid (JA) treatment. Different letters above the box plots indicate statistically significant differences ($p < 0.05$) based on Tukey's HSD test. SolA concentrations are expressed as picomoles per gram of fresh root weight (pmol/g FRW).

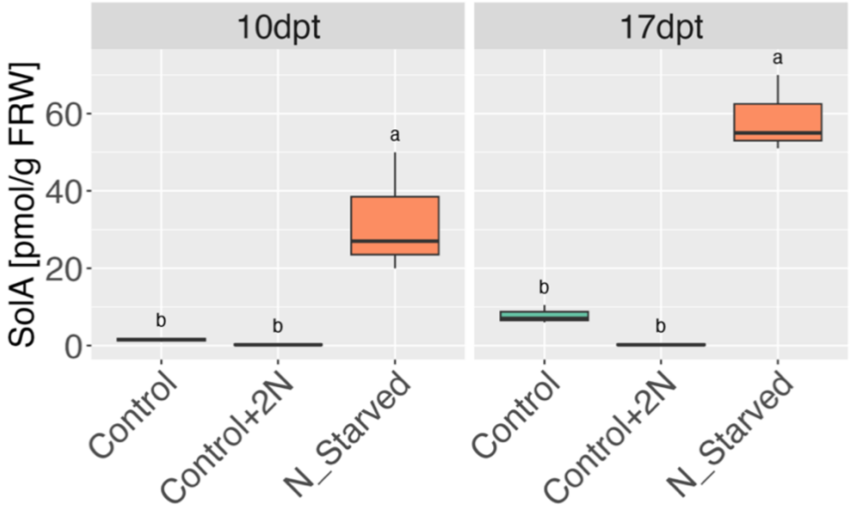


**Figure S2**. SolA concentration in root exudates of tomato plants under nitrogen free (N_Starved), normal nitrogen (Control) and twice the standard nitrogen level (Control+2N) conditions. SolA concentrations are expressed as picomoles per gram of fresh root weight (pmol/g FRW and the sampling occurred after 10 and 17 days post treatment (dpt). Different letters above the box plots indicate statistically significant differences ($p < 0.05$) based on Tukey's HSD test.

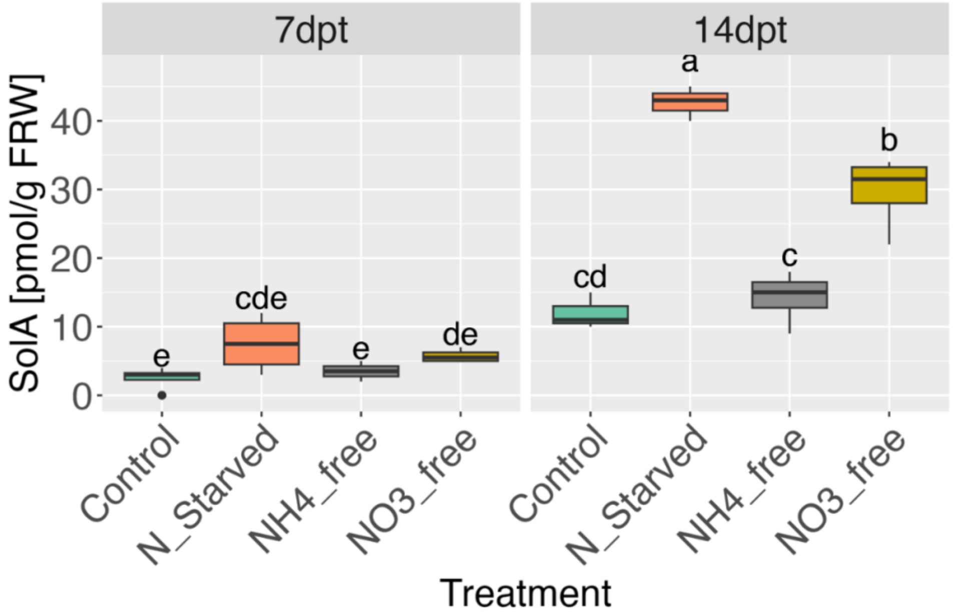


**Figure S3**. Effects of nitrogen conditions on SolA accumulation in tomato root exudates. Tomato plants were cultivated under four nitrogen conditions: nitrogen-starved (N_Starved), standard nitrogen (Control), ammonia-deficient (NH4_free), and nitrate-deficient (NO3_free). SolA concentrations are expressed as picomoles per gram of fresh root weight (pmol/g FRW). Different letters above the box plots indicate statistically significant differences ($p < 0.05$) based on Tukey's HSD test.

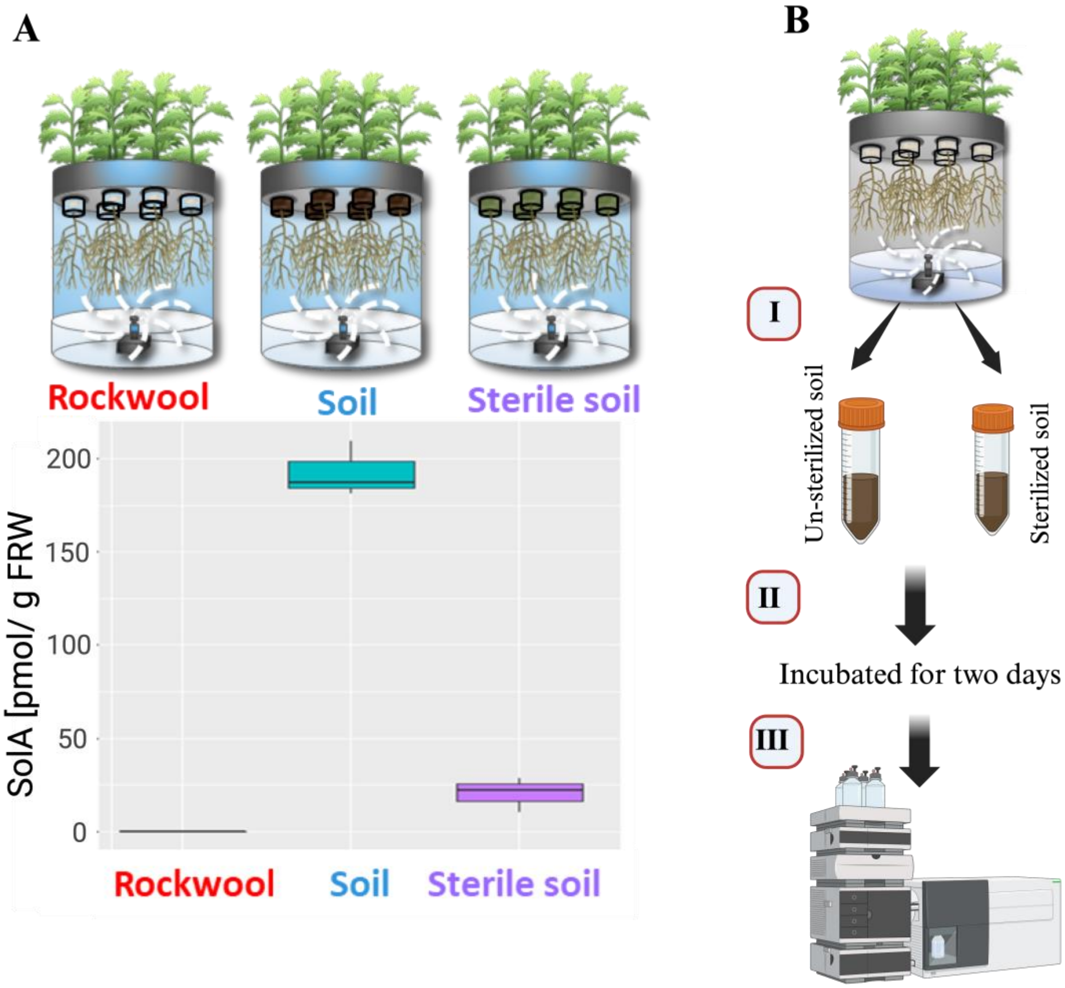


**Figure S4.** A) Comparison of SolA production in tomato plants grown on rockwool, soil, and sterile soil substrates using aeroponic systems. B) Experimental workflow for the conversion assay. Root exudates were collected from nitrogen starved tomato plants growing on rockwool substrate (I) and incubated with unsterilized and sterilized MiCrop soil for two days at room temperature (II), then SolA was extracted and quantified using LC-MS/MS (III). SolA concentrations are expressed as picomoles per gram of fresh root weight (pmol/g FRW).

**Table S1**. Metabolic features associated with SolA biosynthesis identified through the conversion assay using LC-QTOF-MS. Root exudates (RE) were collected from nitrogen-starved tomato plants grown on a rockwool substrate and incubated with both normal and sterilized soils for two days. The table presents m/z values in positive mode, molecular exact mass, retention times (RT), and predicted molecular formulas determined using SmartFormula. The average peak area from five samples of normal soil (AvSoil) and sterile soil (AvS_Soil), along with their ratio (Soil/S_Soil), are reported. A t-test was performed to assess statistical differences between these two groups.

| RT [min] | m/z | M | Molecular Formula | AvSoil | AvS_Soil | Soil/S_Soil | T-TEST |
|---|---|---|---|---|---|---|---|
| **7.87** | 485.1804 | 484.1731 | C26H28O9 | 460.8 | 24.4 | 18.89 | 0.0015 |
| **7.34** | 501.2108 | 500.2035 | C27H32O9 | 411.2 | 78.0 | 5.27 | 0.0487 |
| **6.75** | 499.1944 | 498.1871 | C27H30O9 | 307.2 | 76.4 | 4.02 | 0.0039 |
| **7.04** | 487.1959 | 486.1886 | C26H30O9 | 444.0 | 191.6 | 2.32 | 0.0671 |
| **8.79** | 489.2112 | 488.2039 | C26H32O9 | 318.4 | 181.6 | 1.75 | 0.4728 |
| **8.90** | 459.2371 | 458.2298 | C26H34O7 | 779.2 | 510.4 | 1.53 | 0.2410 |
| **9.36** | 473.2533 | 472.2460 | C27H36O7 | 680.8 | 448.4 | 1.52 | 0.2518 |
| **8.81** | 457.2219 | 456.2146 | C26H32O7 | 417.6 | 287.2 | 1.45 | 0.5278 |
| **7.40** | 473.2175 | 472.2102 | C26H32O8 | 415.6 | 300.4 | 1.38 | 0.6165 |
| **9.02** | 501.2482 | 500.2409 | C28H36O8 | 531.2 | 410.8 | 1.29 | 0.6480 |
| **9.48** | 471.2359 | 470.2286 | C27H34O7 | 358.8 | 318.8 | 1.13 | 0.8314 |
| **8.52** | 543.2226 | 542.2154 | C29H34O10 | 243.2 | 254.8 | 0.95 | 0.9407 |
| **9.01** | 531.2956 | 530.2883 | C30H42O8 | 407.6 | 593.2 | 0.69 | 0.5085 |
| **12.26** | 515.2992 | 514.2919 | C30H42O7 | 176.4 | 308.0 | 0.57 | 0.3195 |
| **11.41** | 499.3049 | 498.2976 | C30H42O6 | 202.8 | 409.2 | 0.50 | 0.1993 |
| **8.78** | 515.2638 | 514.2566 | C29H38O8 | 228.8 | 464.0 | 0.49 | 0.2242 |
| **10.04** | 503.3360 | 502.3287 | C30H46O6 | 105.2 | 815.6 | 0.13 | 0.0117 |

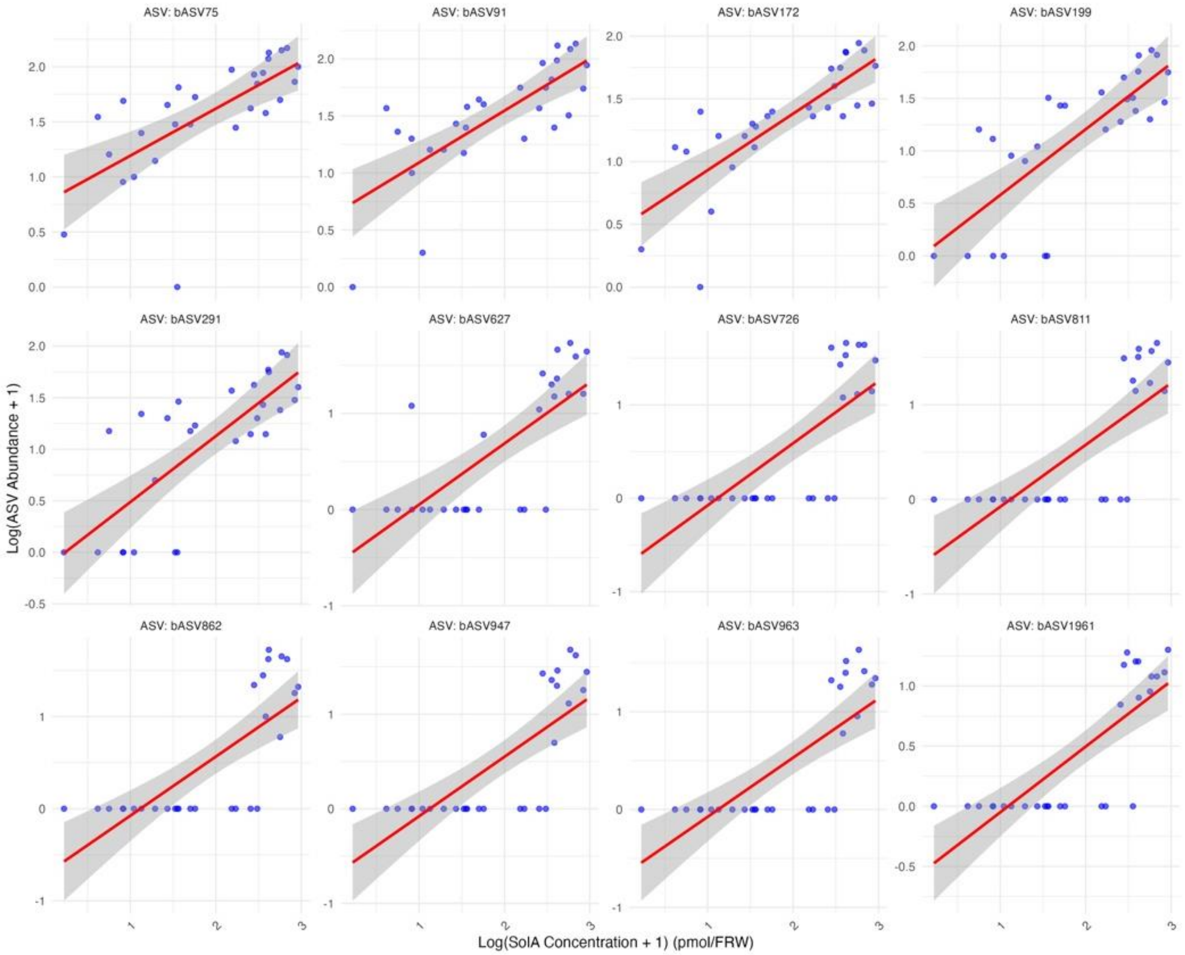


**Figure S5.** Scatter plots illustrate correlation between top 12 bacterial ASVs and SolA concentrations in the rhizosphere compartment. The taxonomical classifications, Spearman correlation coefficients, and p-values of the ASVs are presented in Table S2.

**Table. S2.** Top 12 ASVs positively correlated with SolA content. Spearman correlation coefficients (ρ) and p-values were calculated to assess the relationship between ASV abundance and SolA content. ASVs are ranked in descending order of their correlation coefficient values.

| ASVs | Phylum | Class | Order | Family | Genus | Correlation | p_value |
|---|---|---|---|---|---|---|---|
| **bASV963** | Proteobacteria | Gammaproteobacteria | Burkholderiales | Oxalobacteraceae | Massilia | 0.8128342 | 1.48E-07 |
| **bASV172** | Proteobacteria | Gammaproteobacteria | Burkholderiales | Oxalobacteraceae | Massilia | 0.8827403 | 5.14E-10 |
| **bASV199** | Proteobacteria | Gammaproteobacteria | Burkholderiales | Oxalobacteraceae | Massilia | 0.8107401 | 1.69E-07 |
| **bASV947** | Proteobacteria | Gammaproteobacteria | Burkholderiales | Oxalobacteraceae | Massilia | 0.8102802 | 1.74E-07 |
| **bASV811** | Proteobacteria | Gammaproteobacteria | Burkholderiales | Oxalobacteraceae | Massilia | 0.7995742 | 3.32E-07 |
| **bASV726** | Proteobacteria | Gammaproteobacteria | Burkholderiales | Oxalobacteraceae | Massilia | 0.7986163 | 3.52E-07 |
| **bASV1961** | Proteobacteria | Alphaproteobacteria | Rhizobiales | Xanthobacteraceae | Afipia | 0.7971119 | 3.84E-07 |
| **bASV862** | Proteobacteria | Gammaproteobacteria | Burkholderiales | Oxalobacteraceae | Massilia | 0.7938265 | 4.63E-07 |
| **bASV75** | Proteobacteria | Gammaproteobacteria | Burkholderiales | Oxalobacteraceae | Massilia | 0.7863692 | 7.03E-07 |
| **bASV627** | Proteobacteria | Gammaproteobacteria | Burkholderiales | Oxalobacteraceae | Massilia | 0.7826122 | 8.61E-07 |
| **bASV91** | Proteobacteria | Gammaproteobacteria | Burkholderiales | Oxalobacteraceae | Massilia | 0.7728333 | 1.44E-06 |
| **bASV291** | Proteobacteria | Gammaproteobacteria | Burkholderiales | Oxalobacteraceae | Massilia | 0.7719941 | 1.50E-06 |

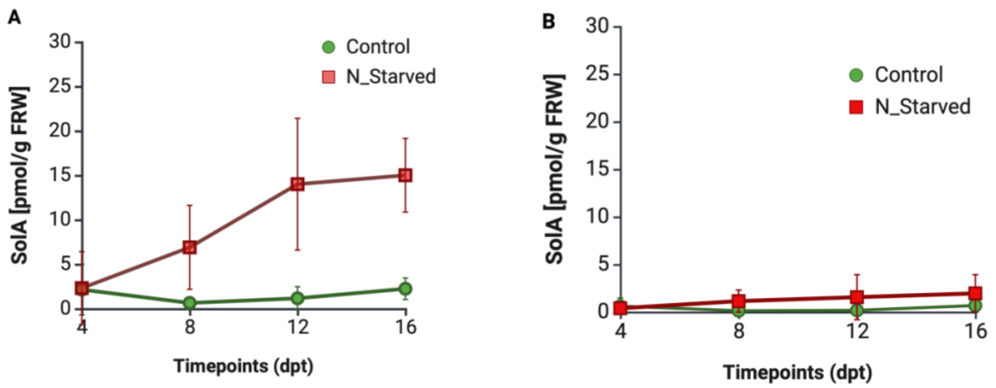


**Figure S6**. Line graph showing SolA concentration in tomato root exudates over time (4, 8, 12, and 16 days post-treatment). Plants were grown on either soil (A) or rockwool (B) and subjected to nitrogen deficiency (red line) or normal nitrogen conditions (green line). SolA concentration is normalized by gram fresh root weight (pmol/g FRW).

**Table S3**. PERMANOVA results for RNA-seq data, showing the effects of day, substrate, and growth condition on the variation in gene expression. Degrees of freedom (df), sum of squares (Ssq), **proportion of explained variance** ($R^2$), F-statistics (F), and **p-value** (Pr(>F)) are provided. Residual variance represents unexplained variation.

| Term | df | Ssq | R2 | F | Pr(>F) |
|---|---|---|---|---|---|
| **Timepoints** | 3 | 41617.11 | 0.164127 | 26.3016 | 0.001 |
| **Substrate** | 2 | 25069.06 | 0.098866 | 23.76509 | 0.001 |
| **Growth condition** | 1 | 86622.35 | 0.341616 | 164.2334 | 0.001 |
| **Residual** | 95 | 49578.84 | 0.195526 | | |

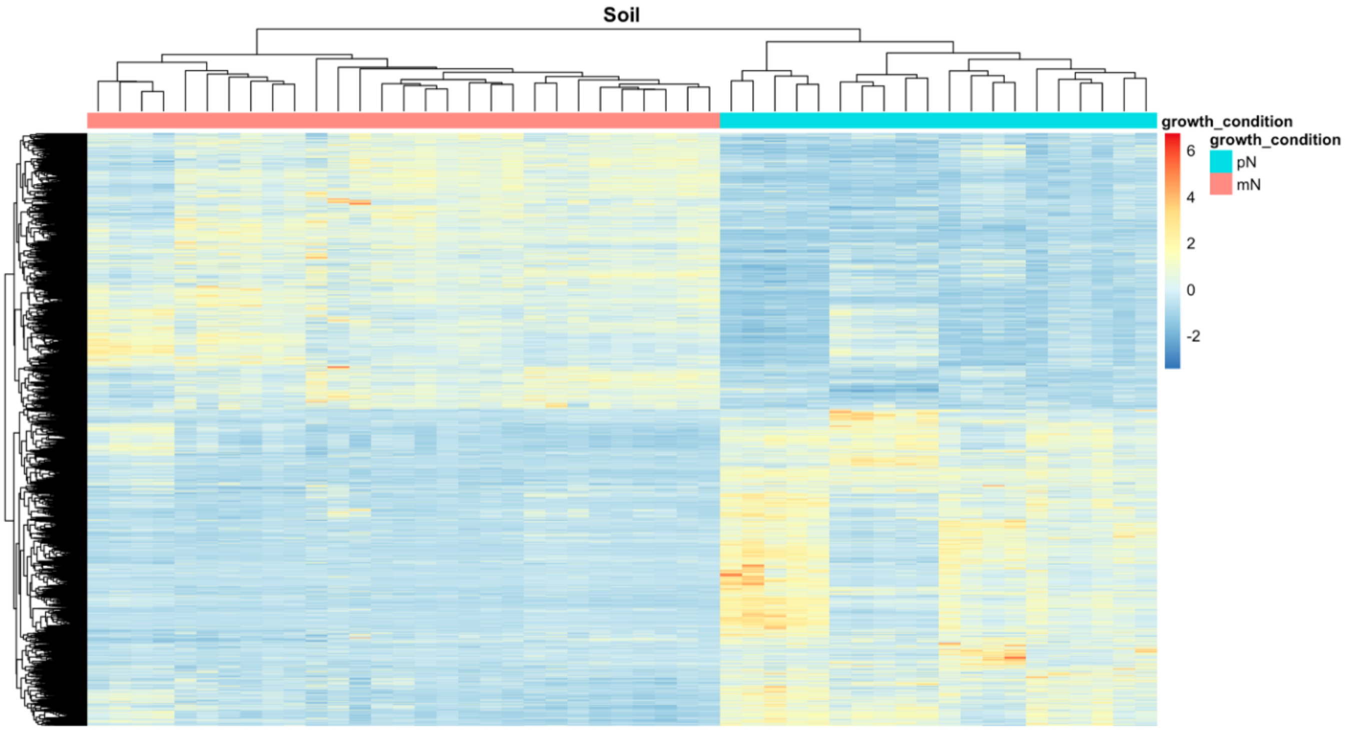


**Figure S7.** Heatmap illustrating gene expression profiles in tomato plants under different nitrogen conditions in soil substrate. The comparison highlights upregulated and downregulated genes between nitrogen-deficiency (mN) and normal nitrogen (pN) treatments in all timepoints, using using a cutoff of |log2FoldChange| > 1 and *p*-value < 0.001. The clustering pattern indicates distinct transcriptional responses influenced by nitrogen availability, with rows representing individual genes and columns representing biological replicates.

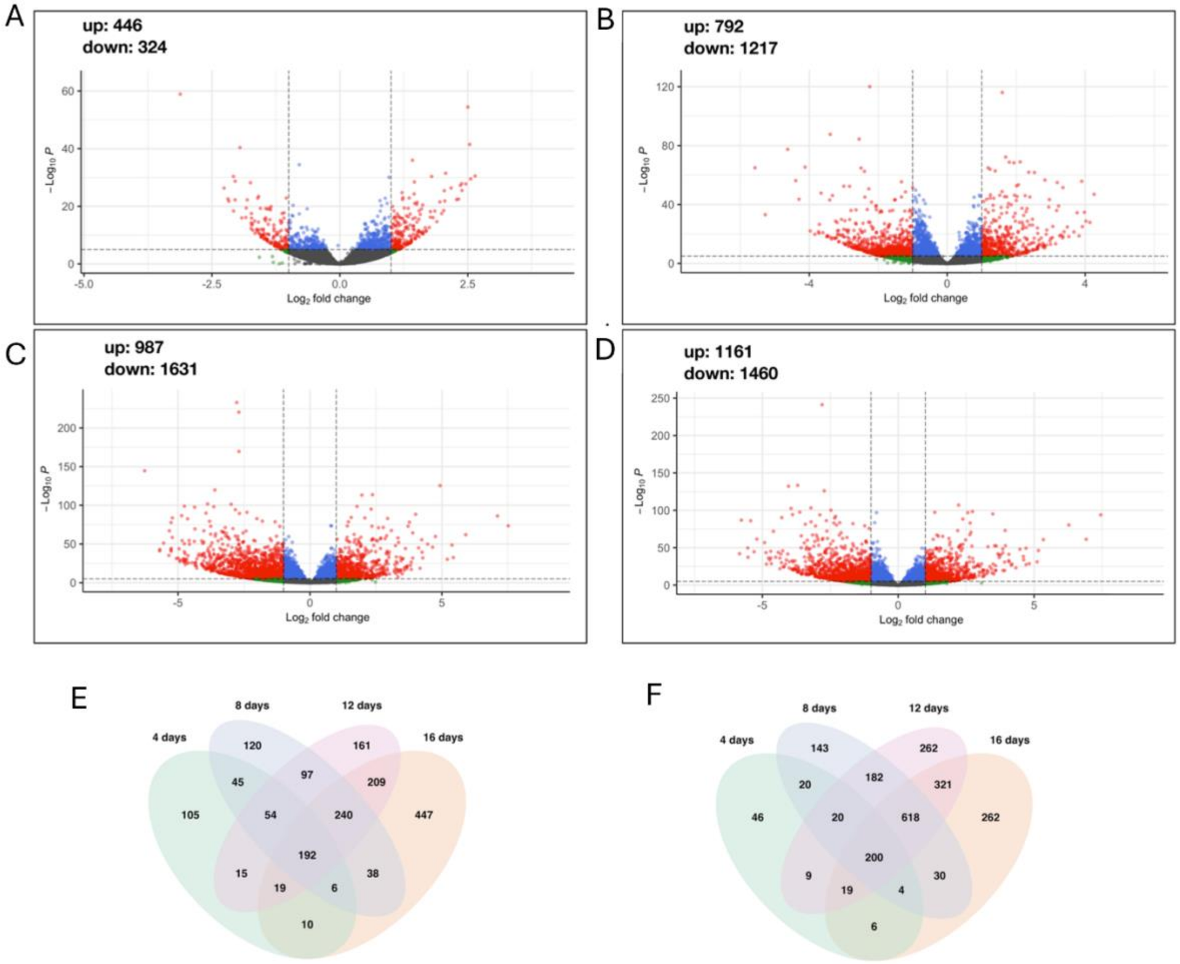


**Figure S8.** Volcano plots and Venn diagrams showing differently expressed genes (DEGs) between plants grown at different timepoints under N deficiency and the respective controls, regardless of the substrate. Volcano plots of 4 days (A), 8 days (B), 12 days (C) and 16 days (D) of N deficiency treatment. With indication of up-regulated (up) and down-regulated (down) DEGs.

Venn diagrams of up-regulated (E) and down-regulated (F) DEGs at different timepoints.

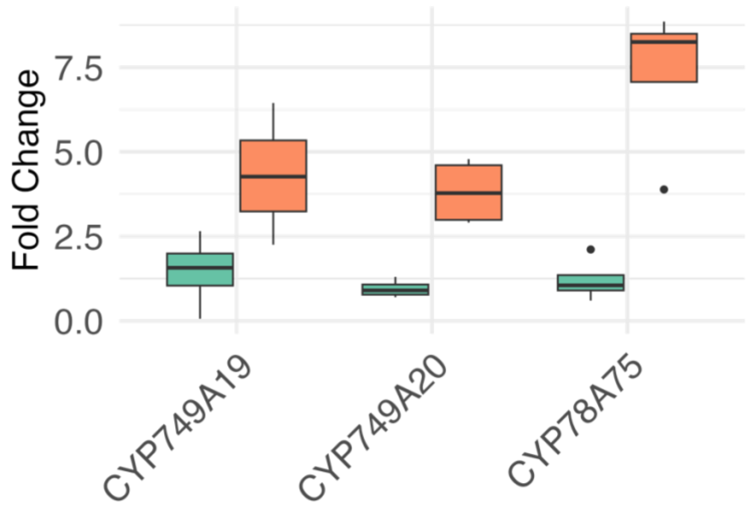


**Figure S9**. Boxplot illustrating the fold change in expression of *CYP749A19*, *CYP749A20*, and *CYP78A75* genes in tomato roots under nitrogen sufficient (Control) and nitrogen starved (N_Starved) conditions. Different letters above the box plots indicate statistically significant differences ($p < 0.05$) based on Tukey's HSD test.

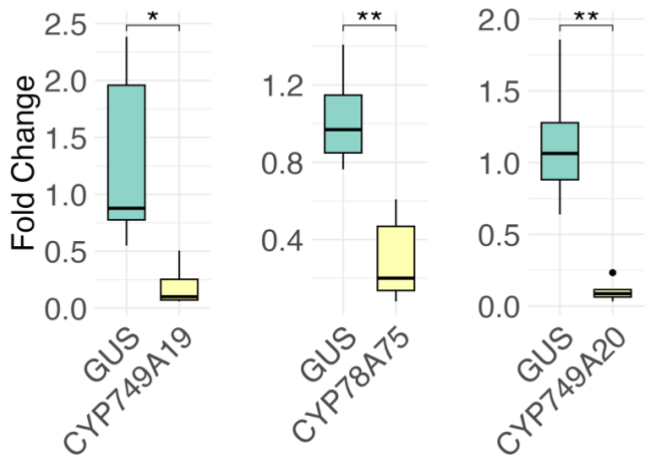


**Figure S10.** Boxplot showing the relative fold change in expression of *CYP749A19* (*Solyc05g011940*), *CYP78A75* (*Solyc03g114940*), and *CYP749A20* (*Solyc05g011970*) in tomato roots following VIGS treatment. Each box represents the fold change of the silenced gene compared to the control (*GUS*). Gene expression levels were normalized to a reference gene, and fold changes were calculated relative to the control. Asterisks indicate statistically significant differences between groups (*$p < 0.05$; $p < 0.01$).

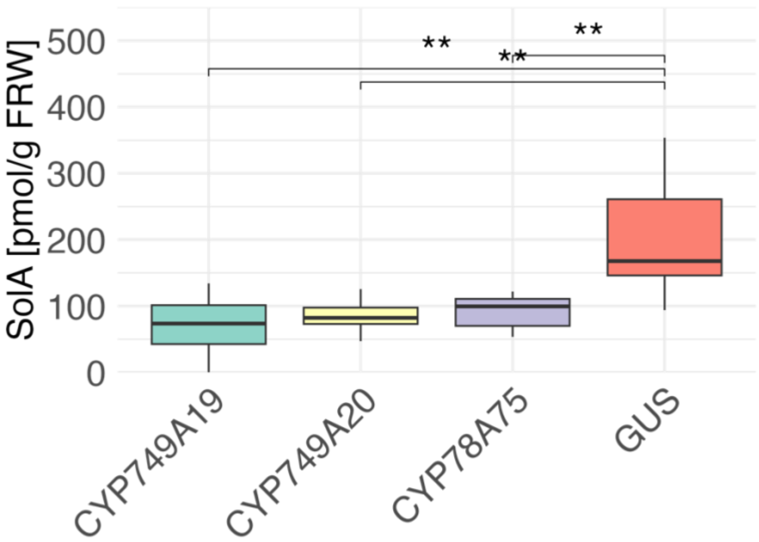


**Figure S11**. SolA concentration in picomole corrected by fresh root weight (pmol/g FRW) in tomato root exudates for plants with silenced genes using VIGS. The x-axis represents the silenced genes (*CYP749A19*, *CYP749A20* and *CYP78A75*) and *GUS* as control. Statistical significance between treatments is indicated by stars (*$p < 0.05$, **$p < 0.01$).

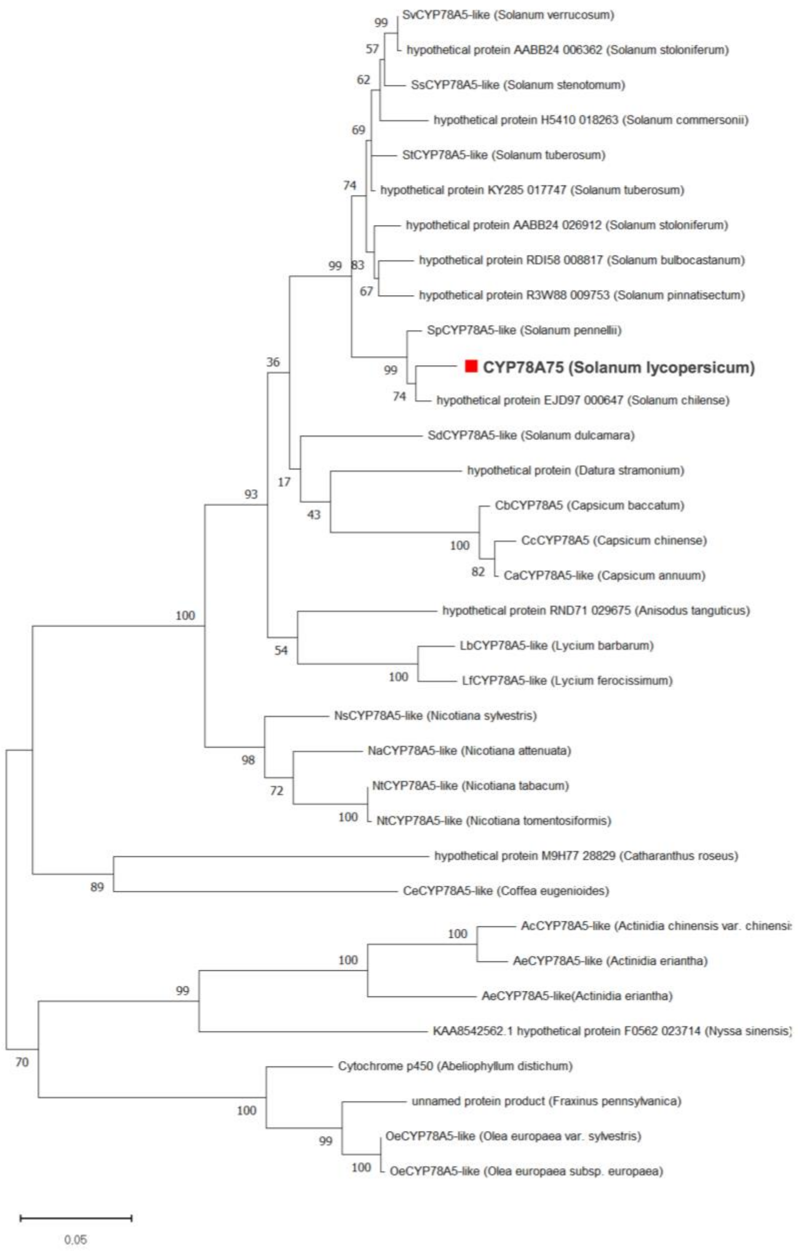


**Figure S12.** Phylogenetic tree for CYP78A75 (marked with a red box) and homologues amino acid sequences constructed using the Maximum likelihood method with 1000 bootstrap replicates.

**Table S4. The list of primers used in this study.**

| Primer ID | Forward | Reverse | Amplicon size | Experiment |
|---|---|---|---|---|
| **CYP749A19** | AACCGCGAACATCAATCACT | CCATGGCTTTGTTTGTGGAC | 219 | VIGS |
| **CYP749A20** | TGGATGCCACTTCAAACACA | CATCCTTGTCCATGTGTGAACA | 189 | VIGS |
| **CYP78A75** | CACACCTCCACTACCACCTT | AAGACCAAGCAAGACCACCT | 230 | VIGS |
| **RT_ CYP749A19** | AAGGTCCTTGTTACAAGTTCCT | AGTTCAGGTTGCAGTCCATG | 187 | qRT-PCR |
| **RT_ CYP749A20** | TTGCTTCTATCTCCTATGGACAC | GCCTTTAATGCCCTGAGAGT | 100 | qRT-PCR |
| **RT_ CYP78A75** | TCCTTCAACTGTTGTGCTCA | ATGGAGACGCGAGCTTTAGA | 123 | qRT-PCR |
| **attB sites** | GGGGACAAGTTTGTACAAAAAA GCAGGCT | GGGGACCACTTTGTACAAGAAA GCTGGGT | - | Gateway cloning |
| **16S_ONT** | AGAGTTTGATCMTGGCTCAG | CGGTTACCTTGTTACGACTT | 1500 | 16S rDNA sequencing |

**Table S5.** Gene IDs, homologous entries in the tomato (solgenomics) and potato genome databases (Spu DB), and the corresponding target regions (including attB and primers) for SolA knocked down genes in tomato via Virus-Induced Gene Silencing.

| Gene ID | Tomato database | Potato database | Target region |
|---|---|---|---|
| **CYP749A19** | Solyc05g011940 | Soltu.DM.05G001230 | TCATAACCTTGAACTCTCAAGTCCATTCAATATCTT GATTCAAAAAAAAAAAAACCGCGAACATCAATCAC TAAGATGATGATGATCATAATCATTCTCGCGAGTT CTCTCTTCATTCTCGTTGGGATCGTTAAGAAGCTA TTATGGACTCCATTTCATGTTCAATTTATGATGAGA TATCAAGGTATACAAGGTCCTTGTTACAAGTTCCTA TATGGGAACTTCAAAGAGATTGATGAAATGAAAAA AGAGTCCACAAACAAAGCCATGGATCACTTATCAC ATGACATATTTCCAAGA |
| **CYP749A20** | Solyc05g011970 | Soltu.DM.05G001210 | ATGGATATAGGGAAACTTATCTTAGTCTTCTTTTTC AGTACTCTTTGCTTCTATCTCCTATGGACACTCTTA AAATTCGTTTATTCAGTATGGTGGATGCCACTTCAA ACACAAAATAAAATGAACTCTCAGGGCATTAAAGG CCCTTCTTATAGTTTTCCTCATGGAAATACCAAAGA TATATCACTGATGAGAAGTCAAACTATGGATAAAC CTATGATTGATATTTCTCATGACATTTTTTCAAGGA TTCAACCTCATGTTCACACATGGACAAGGATGTAC GGGAGGAATTTCCTC |
| **CYP78A75** | Solyc03g114940 | Soltu.DM.03G029090.2 | TGGTCTTCTCCACACCTCCACTACCACCTTCTCCT CTCTCTCTCCAAAACAGAATTCACCTCTTAACTACT CGGCACCTGCATAAACTAAGCAATCTTTGAAGCTC AGAAATGTATCCTGAATACTCTCTTCTCTTCATTCC TTCAACTGTTGTGCTCAACTTAGAGCTTCTTCTCTT TTTTCTCCTCTTTTTTGCGGTTTTCGCCTTCTGGCT TACCCCAGGTGGTCTTGCTTGGTCTTTTTCTAAAG CTCGCGTCTCCATTCCTGGACCTTCTGGTTTACCT CTTCTTGGGTTGGTCT |

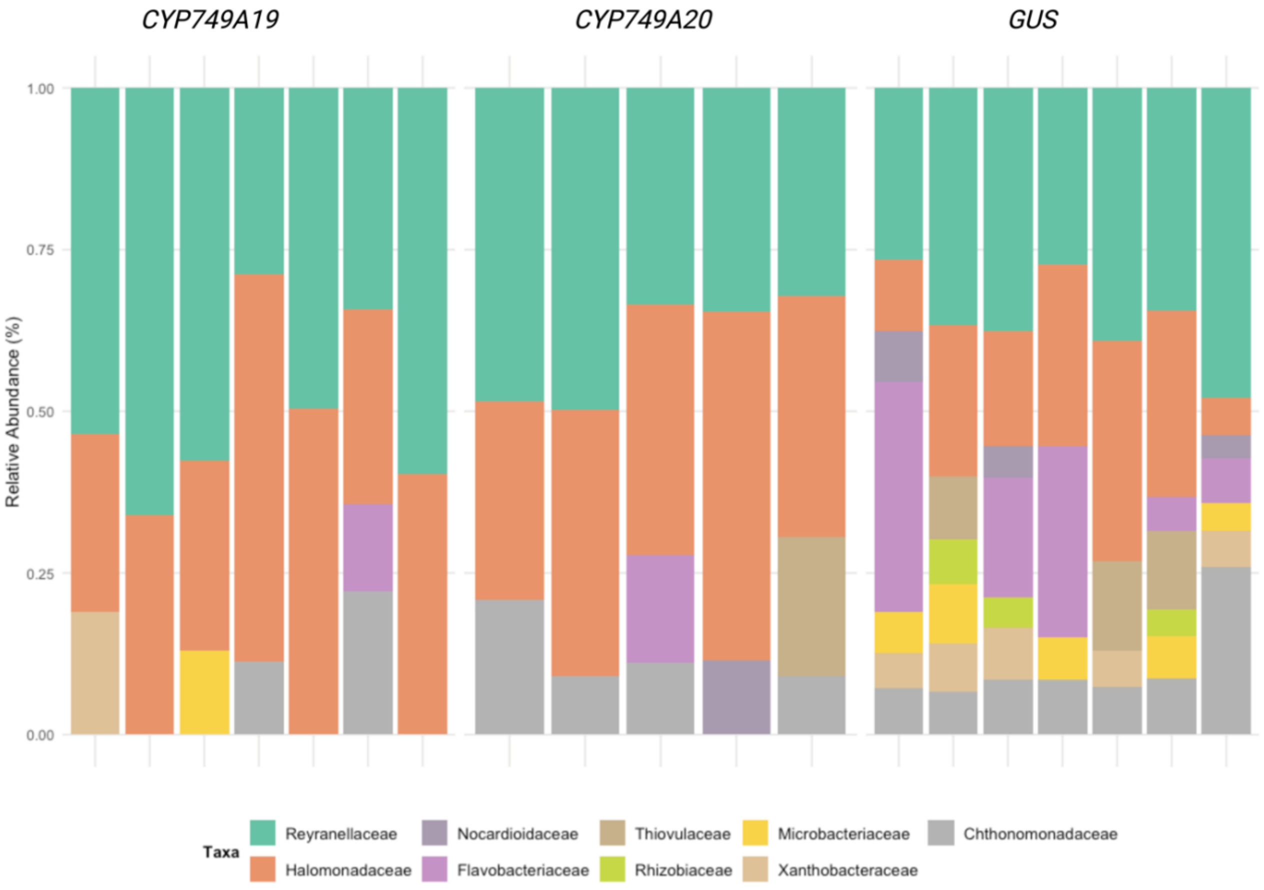


**Figure S13.** Relative abundance of selected microbial taxa in the tomato rhizosphere across silenced plants (*CYP749A19*, *CYP749A20*) and GUS control. These taxa were identified based on comparative analysis between silenced plants and the control, highlighting changes in rhizosphere microbiome composition associated with gene silencing.

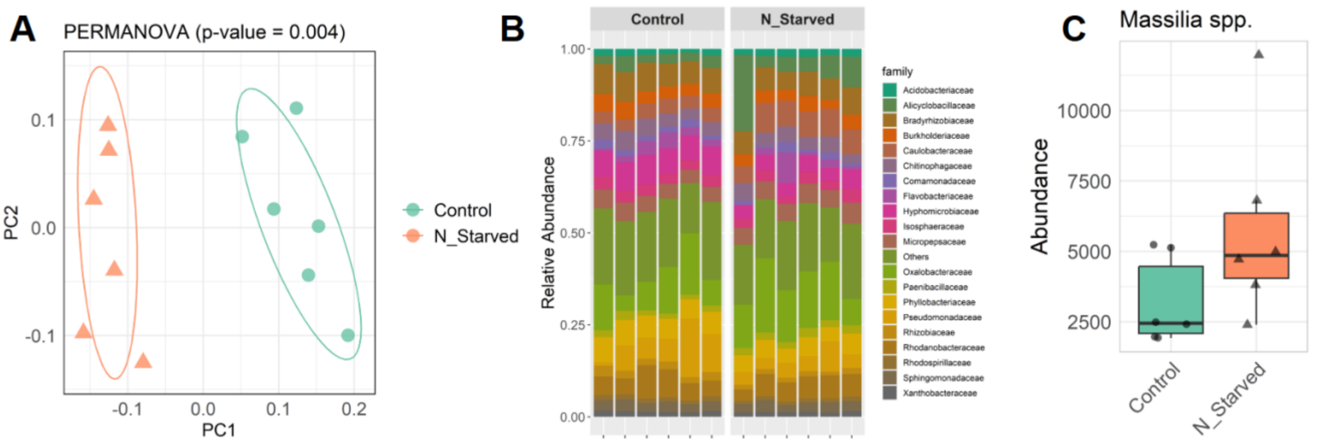


**Figure S14.** (A) Rhizosphere bacterial community changes in tomato plants grown under nitrogen-sufficient (Control) and nitrogen-deficient (N_Starved) conditions after two weeks of nitrogen starvation. A) Principal coordinate analysis (PCoA) shows significant separation of microbial communities between treatments. (B) Relative abundance of bacterial families across samples. (C) Abundance of *Massilia* spp., in the rhizosphere of nitrogen-sufficient (Control) and nitrogen-deficient (N_Starved) tomato plants.

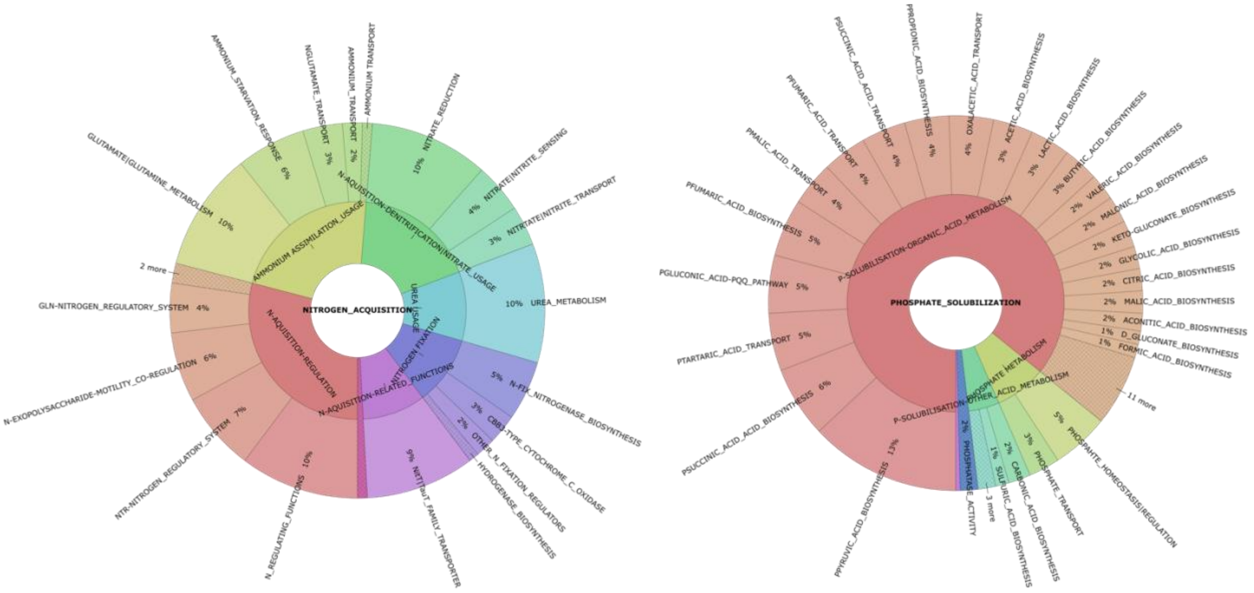


**Figure. S15.** Krona plots depicting nitrogen- and phosphate-related genes identified in the *Massilia* sp. GER05 genome using PlaBase database. Inner rings represent major functional categories including nitrogen fixation/acquisition and phosphate solubilization, while outer rings detail specific genes contributing to these processes.

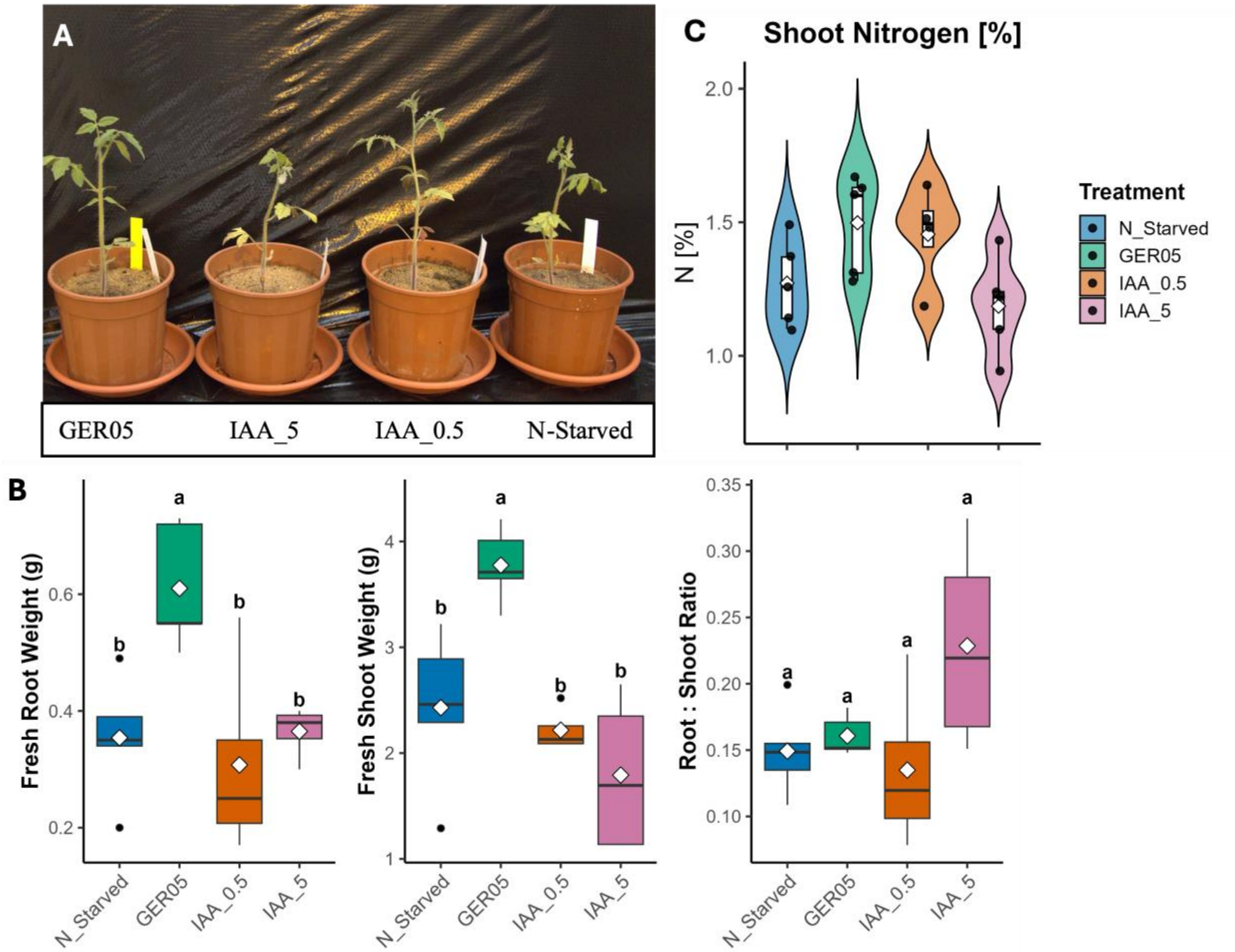


**Figure S16.** Representative tomato plants and effects of nitrogen starvation, *Massilia* sp. GER05 inoculation, and indole-3-acetic acid (IAA) application on biomass and shoot nitrogen content. (A) Representative plants grown under the four treatments: nitrogen starvation (N_Starved), co-inoculation with *Massilia* sp. GER05 (GER05), and supplementation with 0.5 mg/L or 5 mg/L IAA (IAA_0.5 and IAA_5, respectively). (B) Fresh root weight, shoot weight, and root: shoot ratio. (C) Shoot nitrogen concentration (% dry weight). Data are presented as boxplots (median, interquartile range, whiskers 1.5× IQR, white diamonds = mean), with violin distributions and individual data points overlaid where applicable. Statistical differences were performed by one-way ANOVA followed by Tukey's HSD post hoc test. Different letters above boxplots indicate statistically significant differences between treatments ($p$ < 0.05).

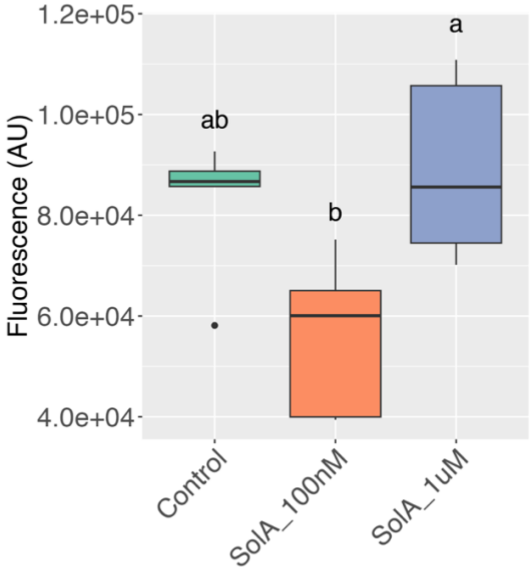


**Figure S17.** Chemotaxis assay of *Massilia* sp. GER05 towards pure SolA with two concentrations (100 nM and 1 μM) and control (milli-Q water).

**Material and Methods**

**Conditions affecting SolA production**

Potato tubers (*Solanum tuberosum* L.) were grown under controlled conditions using an aeroponic system and maintained for three weeks in ¼ Hoagland solution. For salicylic acid (SA) treatments, a stock concentration of 1 M was prepared by dissolving SA in absolute ethanol, from which 1.5, 0.75, and 0.15 mL was added to 3 L of Hoagland solution to achieve final concentrations of 500 μM, 250 μM, and 50 μM, respectively. Control plants were treated with 0.1% ethanol. After one week treatment, 6 ml of the remaining Hoagland solution was collected for SolA extraction. The volume of remaining Hoagland solution was measured, and hanging roots were collected and weighed for SolA quantification.

For high temperature treatment, potato plants were grown in a greenhouse at 22 degrees for three weeks; afterwards, a group of plants were transferred to another compartment and grown for 10 days at 28 degrees. To assess the impact of drought stress on SolA production, two-week-old potato plants were subjected to a one-week period of water deprivation, while control plants continued to receive regular irrigation using demineralized water. For jasmonic acid (JA) treatment, 500 mg of JA was dissolved in ethanol to create a stock solution. A working solution was prepared by diluting 25 μL of the JA stock solution into 50 mL of mili-Q water. Similar to SA, control plants were treated with 50 mL of mili-Q water containing the equivalent volume of ethanol without JA to account for solvent effects. The JA treatment was applied for a duration of one week as well. To collect exudates from the abovementioned potato plants, each pot was flushed using demineralized water. Immediately, 200 mL of the flow-through was collected, from which a 6 mL aliquot was taken for SolA extraction. Moreover, plant roots were carefully washed and weighed for SolA quantification.

**PCN hatching assay**

The hatching experiment was conducted using *Globodera pallida* D383 (pathotype Pa3), which was propagated on Solanum tuberosum and stored at 4°C. Briefly, cysts were initially soaked in tap water and incubated at 20°C in darkness for seven days. Following incubation, the cysts were carefully crushed and sieved to collect the eggs. A 100 µL suspension containing approximately 100 eggs was transferred into glass-coated plates. Root exudates collected from Hoagland solution of nitrogen-starved and nitrogen-sufficient plants were filtered through a 0.22 µM filter, diluted tenfold, and added to each well. In this experiment, root exudates from the high SolA-producing potato variety MFII were used as a positive control, while only Hoagland solution and tap water served as negative controls.

The plates were maintained at 20°C in darkness for ten days. Photographs of each well were taken with a stereomicroscope at the beginning and end of the assay. The numbers of hatched juveniles and unhatched eggs were counted, and the hatching percentage was calculated using the formula:

$$\left[\frac{Jt10 - Jt0}{Et0}\right] \times 100$$

where Jt10 represents the number of hatched juveniles after 10 days of treatment, Jt0 is the number of spontaneously hatched juveniles at the beginning of the experiment, and Et0 is the total eggs at the start of the assay.

**Soil collection, plant growth and microbiome analysis**

To evaluate the impact of soil type on SolA production, tomato plants (*Solanum lycopersicum* cv. Money maker) were grown in the pots filled with various collected soils. Different potting soils (Type I, II and III) were sourced from breeding companies in the Netherlands, while forest soil was collected from a natural heathland in Hoge Veluwe National Park, the Netherlands. The plants were kept at 22°C for four weeks and irrigated with tap water. To collect root exudates, each pot was flushed using demineralized water. Immediately, 200 mL of the flow-through was collected, from which a 6 mL aliquot was taken for SolA extraction. Moreover, plant roots were carefully washed and weighed for SolA quantification.

In a separate experimental setup, forest soil was mixed with various potting soils at a 1:9 ratio (forest soil: potting soil). Similar to previous experiment, tomato plants were grown in these soil mixtures and root exudates were collected from the plants for SolA quantification.

Rhizosphere soil and root compartments were collected for microbiome analysis. To collect rhizosphere soil, tomato plants were carefully uprooted, and loosely adhering soil was gently shaken off, leaving only soil tightly associated with the roots. This rhizosphere soil was then collected by further agitating the roots in a sterile solution to detach the remaining soil particles. Both root and rhizosphere soil samples were then processed for microbial DNA extraction using the PowerSoil DNA Isolation Kit (QIAGEN) according to the manufacturer's instructions. Extracted DNA was subjected to 16S rDNA sequencing on an Illumina MiSeq PE250 platform at Génome Québec Innovation Centre (Montréal, QC, Canada), targeting the V3–V4 region. The sequencing raw data were preprocessed using the DADA2 package integrated with QIIME2 (Quantitative Insights Into Microbial Ecology, version 2) for high-resolution microbial community profiling. Forward and reverse reads were trimmed to 220 bp and 200 bp, respectively. Denoised paired-end reads were merged, and chimeric sequences were identified and removed to enhance accuracy. Amplicon Sequence Variants (ASVs) were assigned to taxonomic identities using the SILVA database (version 138).

After obtaining ASV count table, spearman correlation analysis was performed to evaluate the monotonic relationship between the relative abundance of each ASV and SolA content across samples. Correlation coefficients and corresponding *p*-values were calculated for each ASV and top significant correlations ($p < 0.05$) were used for illustration.

**Phylogenetic tree construction**

The amino acid sequences of the *CYP78A75* gene were retrieved from https://solgenomics.net and used to identify homologous sequences with over 75% similarity through BLASTp. The selected sequences were downloaded in FASTA format, and amino acids sequence alignment was performed using the ClustalW algorithm within the MEGA11 software package. Moreover, the candidate cytochrome P450 monooxygenase (P450) and 2-oxoglutarate/Fe(II)-dependent dioxygenase (ODD) genes identified through co-expression analysis were subjected to phylogenetic analysis to examine their evolutionary relationships. P450 sequences were further classified into subfamilies using the P450 database (https://p450atlas.org/search). Phylogenetic trees for *CYP78A75,* P450s and ODDs were generated independently in MEGA 11 using the Maximum Likelihood method with 1000 bootstrap replicates.

**Tomato nitrogen starvation treatment**

Tomato (*Solanum lycopersicum*) seeds were surface-sterilized and pre-germinated at 24 °C for three days. Pre-germinated seeds were transferred to pots containing a compost: sand mixture (15:85, v/v) and maintained in a greenhouse under controlled conditions (22 °C, 16 h light/8 h dark photoperiod). Plants were supplied with one-quarter strength Hoagland nutrient solution once per week for two weeks.

After two weeks, pots were flushed twice with deionized (Dami) water to remove residual nutrients. Plants were subsequently grown for an additional two weeks under two treatments: (i) nutrient-sufficient conditions (one-quarter strength complete Hoagland solution), and (ii) nitrogen-starved conditions (one-quarter strength Hoagland solution lacking $NH_4NO_3$). Additionally, plants were regularly watered with deionized (Dami) water for optimal growth and to avoid drought stress. Rhizosphere soil collection, microbial DNA extraction, and metabarcoding of 16S rDNA were carried out as described below.

**Microbial DNA extraction and nanopore sequencing of 16S rDNA**

To evaluate the microbiome compositions in the rhizosphere of reduced SolA tomato plants, we profiled the microbial communities associated with tomato plants in which key SolA biosynthetic genes, including *CYP749A20* and *CYP749A19*, were silenced using VIGS. For this, rhizosphere soils were collected from tomato plant roots by gently shaking the roots to remove loosely attached soil, retaining only the soil closely adhering to the roots surface in sterilized milli-Q water. The collected rhizosphere soil was freeze-dried, and 50 mg of the dried soil was used for DNA extraction with the DNeasy PowerSoil pro Kit (QIAGEN), following the manufacturer's protocol. DNA concentration and purity were assessed with a NanoDrop spectrophotometer.

To profile the microbial community in the rhizosphere, the entire 16S rDNA was amplified using 16S primers (Supplementary Table S4) using a LongAmp DNA polymerase (New England Biolab). The PCR product was visualized using 1% agarose gel electrophoresis followed by purification with 0.8× AMPure XP beads (Beckman Coulter). The concentration and quality of these purified PCR products were measured using a Nanodrop spectrophotometer (Thermo Scientific). For library preparation, 200 fmol of each sample was used and library was prepared using Native Barcoding kit (SQK-NBD114.96) as per manufacturer's instructions. The purified library (barcoded and adapter ligated) was quantified using a Qubit fluorometer (Invitrogen) and 20 fmol of final library was used for nanopore sequencing.

Sequencing on minion flow cell (R10.4.1) with a MinION MK1b device was performed for 72 h with min q-score value set as 10. Real time super accurate basecalling was performed using Guppy integrated within MinKNOW software. Following sequencing, demultiplexed fastq files for each sample were combined and quality trimming was performed using Nanofilt (De Coster *et al.*, 2018). Finally, the sequences in each sample were taxonomically classified and abundances were calculated using Emu v3.4.6 software pipeline (Curry *et al.*, 2022) against emu database on a Linux server with following parameter --map-ont, --keep-counts, and --keep-read-assignments. The resulting output for each sample were combined using emu combine-output function with --counts, --split-table options to build final count and taxonomy tables, which were used for diversity and differential microbiome analyses.

**Genome sequencing of *Massilia* sp. GER05 and functional genome analyses**

High molecular weight DNA was extracted from *Massilia sp*. GER05 using the Qiagen Blood and Cell Culture DNA Mini Kit (Qiagen, USA) with a 20/G column. The extracted DNA underwent purification with 1× AmPure beads, and its quality and concentration were measured using a NanoDrop 2000 spectrophotometer (Thermo Fisher Scientific, USA). A genomic library was prepared using the Native Barcoding Kit V14 (Oxford Nanopore Technologies, UK), following the manufacturer's protocol, with 1000 ng of the purified DNA. The library yield was determined using a Qubit 3.0 fluorometer (Invitrogen, Life Technologies), and 10 fmol of the prepared library was loaded onto a Flongle R10.4.1 flow cell. Sequencing was performed on a MinION Mk1b device for 24 hours with real-time super-accurate (SUP) basecalling via the MinKNOW software.

The sequencing data were pre-processed using NanoFilt (De Coster *et al.*, 2018), applying a quality threshold of 10 and filtering out reads shorter than 500 bp. Genome assembly was conducted with the Flye assembler (v2.9.3) in high-quality nanopore mode (--nano-hq) (Kolmogorov *et al.*, 2019). The assembled genome was polished using Medaka (v1.7.2) and subsequently annotated with Prokka (v1.14.6). Genes associated with plant growth-promoting traits were identified using the PGPT-Pred tool available at PlaBAse (v1.02) (Patz *et al.*, 2024, 2021). RhizoSMASH (available at https://git.wur.nl/rhizosmash) analysis to detect catabolic gene clusters in the bacterial genome was performed using annotated assembly as input with default parameters.

**Plant bacteria co-inoculation experiment**

To assess the plant growth-promoting effects of *Massilia* sp. GER05 on tomato plants, co-inoculation experiments were performed in greenhouse conditions. Tomato seeds surface-sterilized seeds were germinated, and the seedlings were incubated in a bacterial suspension for 1 hour. Bacterial suspensions were prepared to an $OD_{600}$ of 0.5 in 1× phosphate-buffered saline (PBS) prior to inoculation. The treated seedlings were transplanted into 250 mL pots containing MiCRop soil and sand mixture (15:85). Additionally, 1 mL of bacterial suspension was added directly to the roots of each seedling, followed by covering the roots with soil. Mock-inoculated seedlings, treated with 1× PBS, served as controls. Each treatment included six pots, each containing a single seedling. Plants were grown for three weeks, with watering as needed and weekly supplementation with Hoagland's solution without $NH_4NO_3$. After three weeks, plants were harvested, and biomass and chlorophyll content in leaves were measured.

In the second experiment, the surface sterilized tomato seeds were germinated on complete Murashige and Skoog (MS) medium for one week. Seedlings were then subjected to root-dip treatments for 1 h in one of the following solutions: indole-3-acetic acid (IAA, 0.5 or 5 mg/L), a suspension of *Massilia* sp. GER05 ($OD_{600}$ = 0.5), or phosphate-buffered saline (PBS) as a negative control. Similar to previous experiment, the seedlings were transferred to 250 mL pots containing a soil–sand mixture (MiCRop soil:sand, 15:85). Plants were maintained under controlled greenhouse conditions for three weeks, with watering as required and weekly supplementation with Hoagland's solution lacking $NH_4NO_3$. At harvest, shoots and roots were separated, and fresh biomass was recorded.

To measure nitrogen content, oven-dried shoot samples (5–10 mg) were finely ground. Aliquots (5–10 mg) were weighed into tin capsules and analyzed for total nitrogen by dry combustion on an elemental CN analyzer (Elementar vario EL/EL cube). Each run included procedural blanks and a 15-point sulfanilic acid calibration (1–3 mg per standard). A subset of biological samples (10%) was analyzed in duplicate to account for variability in sample homogeneity. Nitrogen concentration was reported on a dry-mass basis (% N).

**Chlorophyll content determination**

Chlorophyll a and b contents were determined spectrophotometrically, following established protocols by (Lichtenthaler *et al.*, 1996; Porra and Scheer, 2019). For this end, 200 mg of frozen leaf tissue was homogenized in 5 mL of ice-cold 80% aqueous acetone. The extract was then collected, and the tissue was rinsed with an additional 5 mL of acetone, which was combined with the initial extract. Afterwards, samples were incubated on ice for 2 hours with gentle agitation. Then, the samples were centrifuged at 10,000 rpm for 10 minutes at 4°C. The supernatant was collected, and absorbance was measured at 664 nm and 647 nm using a UV-Vis spectrophotometer. Total chlorophyll concentrations (mg/mL) were calculated using the following formulas:

$$[\text{Chl a}] = 12.25 \times A_{664} - 2.55 \times A_{647} \text{ mg/mL}$$

$$[\text{Chl b}] = 20.31 \times A_{647} - 4.91 \times A_{664} \text{ mg/mL}$$

$$[\text{Chls a + b}] = 17.76 \times A_{647} + 7.34 \times A_{664} \text{ mg/mL}$$

**Data analysis**

All data analyses were conducted using R (v4.1.1) and a variety of specialized packages to facilitate data visualization, statistical testing, and differential expression analysis. Visualization of results was carried out using ggplot2 for creating customized box plots and ComplexHeatmap to generate detailed heatmaps.
